# ADAM interact with large protein complexes to regulate Histone modification, gene expression and splicing

**DOI:** 10.1101/2024.08.18.608474

**Authors:** Ankit Pandey, Helene Cousin, Shiv Kumar, Louis Taylor, Ashmita Chander, Kelsey Coppenrath, Nikko-Ideen Shaidani, Marko Horb, Dominique Alfandari

## Abstract

Cranial neural crest (CNC) cells are key stem cells that contribute to most of the facial structures in vertebrates. ADAM (<u>A D</u>isintegrin <u>A</u>nd <u>M</u>etalloprotease) proteins are essential for the induction and migration of the CNC. We have shown that Adam13 associates with the transcription factor Arid3a to regulate gene expression. Here we show that Adam13 modulates Histone modifications in the CNC. We show that Arid3a binding to the *tfap2α* promoter depends on the presence of Adam13. This association promotes the expression of one *tfap2α* variant expressed in the CNC that uniquely activates the expression of gene critical for CNC migration. We show that both Adam13 and human ADAM9 associate with proteins involved in histone modification and RNA splicing, a function critically affected by the loss of Adam13. We propose that ADAMs may act as extracellular sensors to modulate chromatin availability, leading to changes in gene expression and splicing.

## Introduction

Cranial Neural crest (CNC) cells are a transient population of stem cells induced during early vertebrate embryogenesis. They migrate from the lateral side of the neural plate and differentiate into multiple cell types, including those that give rise to most of the cranial skeleton and peripheral nervous system ^1^. The process of induction and migration involves multiple signaling pathways, including the Wnt, BMP and FGF, as well as multiple chemo attractants and chemorepellents to direct cells into the proper tracks and toward specific structures (Koontz et al 2023). We have previously shown that four ADAM cell surface metalloproteases are involved in both the specification and migration of the CNC ^2–5^. Among these, Adam13 (ADAM33 homologue) cleaves Cadherin-11 and the protocadherin PCNS, releasing a portion of their extracellular domain that promotes cell migration ^4,6^. For Cadherin-11, the fragment interacts with EGFR2 receptor to stimulate AKT phosphorylation independently of its homophylic binding site ^7,8^. In the neural crest, ADAMs have been shown to play a key role in cell adhesion and epithelial to mesenchymal transition by cleaving cadherins such as E-cadherin and cadherin-6B ^9–11^. In addition to this clear role of the proteolytic activity of Adam13, the non-proteolytic functions of ADAMs are also critical. For example, Adam11, a non-proteolytic ADAM, negatively regulates Wnt signaling while positively regulating BMP4 signaling to control CNC cell proliferation and EMT ^2^.

We have previously shown that Adam13 can cleave itself within the cysteine-rich domain ^12^, becoming a substrate for γ-secretase ^13^. The cytoplasmic domain then translocates into the nucleus where it regulates the expression of thousands of genes ^13^. The mechanism by which Adam13 regulate gene expression remains unknown, but several targets regulated by Adam13, including Tfap2α, PCNS and Calpain-8, are essential for CNC migration. Other ADAMs, and MMPs have been found in the nucleus, but their exact function or mode of action remains unknown ^14,15^. In esophageal squamous cell carcinoma, ADAM9 was found associated with chromatin in response to hypoxia. Chromatin immunoprecipitation experiment showed that ADAM9 associated with the promoter of gene repressed during hypoxia ^16^.

Arid3a/dril1/BRIGHT is a transcription factor important for the first cell fate decision in mice ^17^ and is important for mesoderm induction and gastrulation in Xenopus where it regulates TGFβ signaling downstream of Smad ^18^. We have previously shown that Adam13 interacts with Arid3a, to increase the expression of the transcription factor *tfap2α* ^6^, a gene important for CNC induction and maintenance ^19,20^. Loss of Tfap2α leads to craniofacial defects in all vertebrates tested ^19,21–24^. In mice *Tfap2α* KO leads to perinatal lethality with craniofacial and neural tube defects ^25^. In Human, mutations in the *TFAP2α* gene leads to Branchiooculofacial Syndrome ^23^, highlighting Tfap2α functional conservation across species. Arid3a can act as an activator or a repressor of gene transcription ^26^. In Xenopus Arid3a has been shown to activate lhx1 gene expression for nephric tubule regeneration by modulating the chromatin landscape on the gene promoter of *lhx1* by interacting with Kdm4 and removing histone H3K9 di-tri methylation (H3K9me2/3) ^27^. Thus, in Xenopus, Arid3a can activate transcription with Smad co-factors during mesoderm induction and regulate transcription through epigenetic modification during organogenesis. In the neural crest, the Arid3a/Adam13 complex does not work via Smad ^6^, suggesting that it may involve histone modification instead.

Recent studies have shown that chromatin organization is changed during neural crest cell development and that this change is essential for both induction and migration ^28–31^. Significant strides have been made over the last three decades in identifying and understanding gene regulatory networks that regulate CNC ^32^, including key transcription factors that exemplify CNC, such as the SOX9, FOXD3, SNAI1, SNAI2, and TFAP2 family ^32^. Alternative splicing (AS) has also emerged as an important regulator of CNC development, highlighted by the human craniofacial birth defects arising from abnormalities in the splicing regulator genes ^33–38^.

In this study, we show how Adam13 regulates *tfap2α* variants differentially by binding to Arid3a and promoting its binding at the promoter of one variant while inhibiting the expression of another. We also show that Adam13 is involved in regulating histone modifications in CNC globally as well as in the promoter region of *tfap2α*. We also identify a novel role of the Tfap2α variant in the regulation of alternative splicing.

## Results

### Adam13 knockdown leads to increased H3K9me2/3 levels

We have previously shown that Adam13 and Arid3a interact at the plasma membrane and translocate to the nucleus, and this process is important for the expression of the transcription factor *tfap2α* in the CNC ^6^. ARID3A/ARID3B have been shown to regulate gene expression by binding to histone-demethylating enzyme KDM4C and reducing H3K9 trimethylation at target genes in Human Cancer and nephric tubule regeneration in Xenopus ^27,39^. To test whether the loss of Adam13 would result in changes in histone methylation, we performed immunofluorescence (IF) to detect H3K9me2/3 in migrating wild-type CNC (NI) or CNC lacking Adam13 (MO13, Fig.1A). In three independent experiments quantifying the intensity of H3K9 relative to DAPI staining of the DNA, we found an increased overall intensity of H3K9me2/3 in CNC explants lacking Adam13 (Fig.1B). This increase in the H3K9me2/3 signal is not due to a change of Arid3a mRNA or protein expression level in CNC lacking Adam13 (Fig.1C, D and Supplementary Fig.S1B) confirming earlier results ^6^. Given that levels of H3K9me2/3 often vary in concert with H3K4 methylation ^40^, we also tested the presence of these marks in CNC lacking Adam13 (Fig.2). CNC explants were co-stained with an antibody to Snai2 to confirm their CNC identity (Fig.2A). We quantified the levels of H3K4me3 in the nucleus of Snai2 positive cells, and we found that H3K4me3 levels were significantly decreased in CNC lacking Adam13 (Fig.2B).

**Figure 1.**
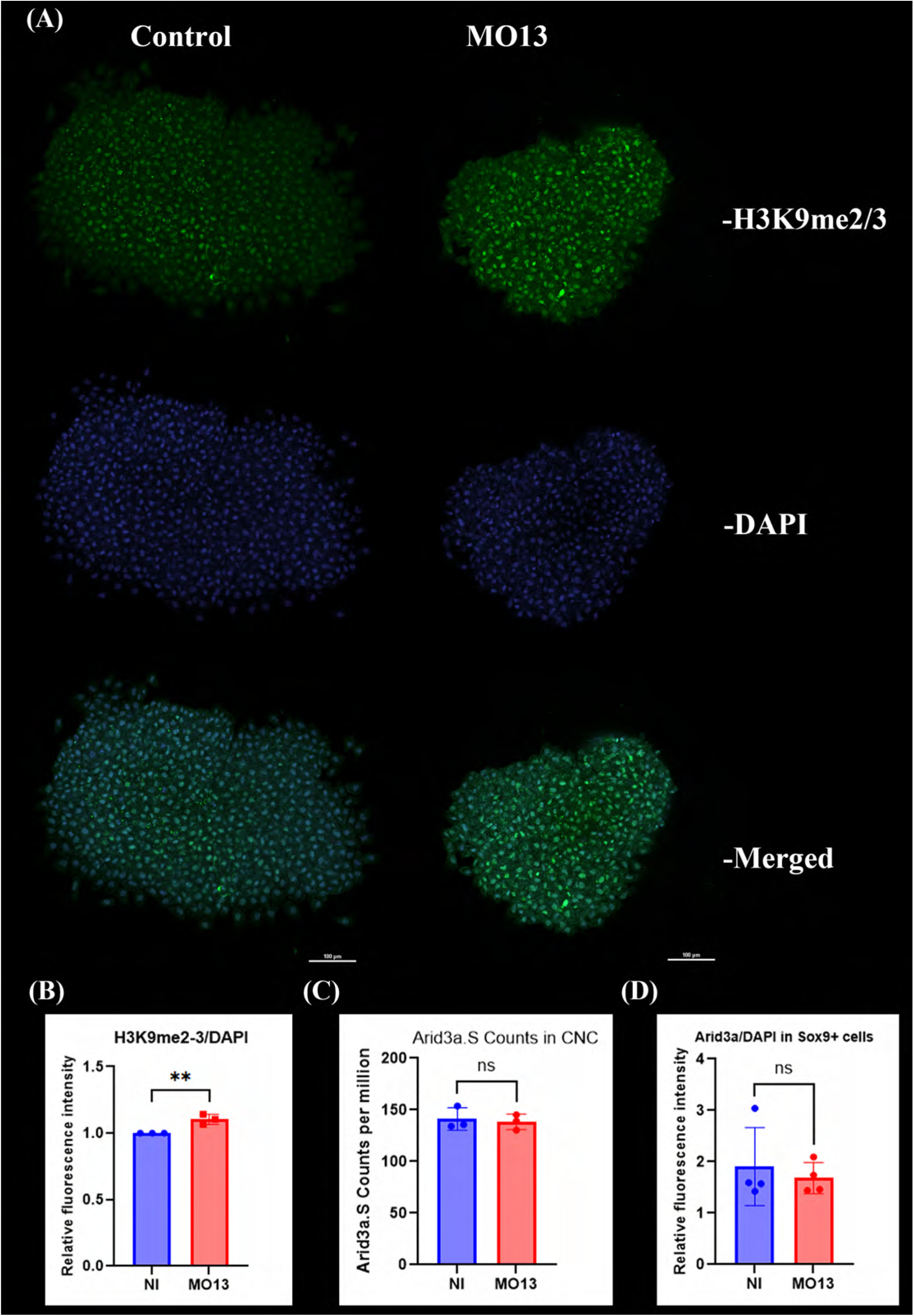
Loss of Adam13 leads to increased H3K9me2/3 methylation in CNC. (A) Immunofluorescence of CNC explants immuno-stained for H3K9me2/3 (green) and DAPI (blue). All images are maximum intensity projection (Max-IP). (B) Histogram depicting the relative fluorescence intensity of H3K9me2/3 between Control (NI) and Adam13KD (MO13) CNC explants. Student’s t-test was performed for statistical analysis, ** p-value<0.01. (C) Histogram depicting counts per million (CPM) of Arid3a.S RNA present in CNC RNA-sequencing. Student’s t-test was performed for statistical analysis following the CPM normalization; not significant (ns) represents p-value>0.05. (D) Histogram depicting the relative fluorescence intensity of Arid3a in Sox9 positive cells between Control (NI) and Adam13KD (MO13) CNC explants. ns represents p-value>0.05.

**Figure 2.**
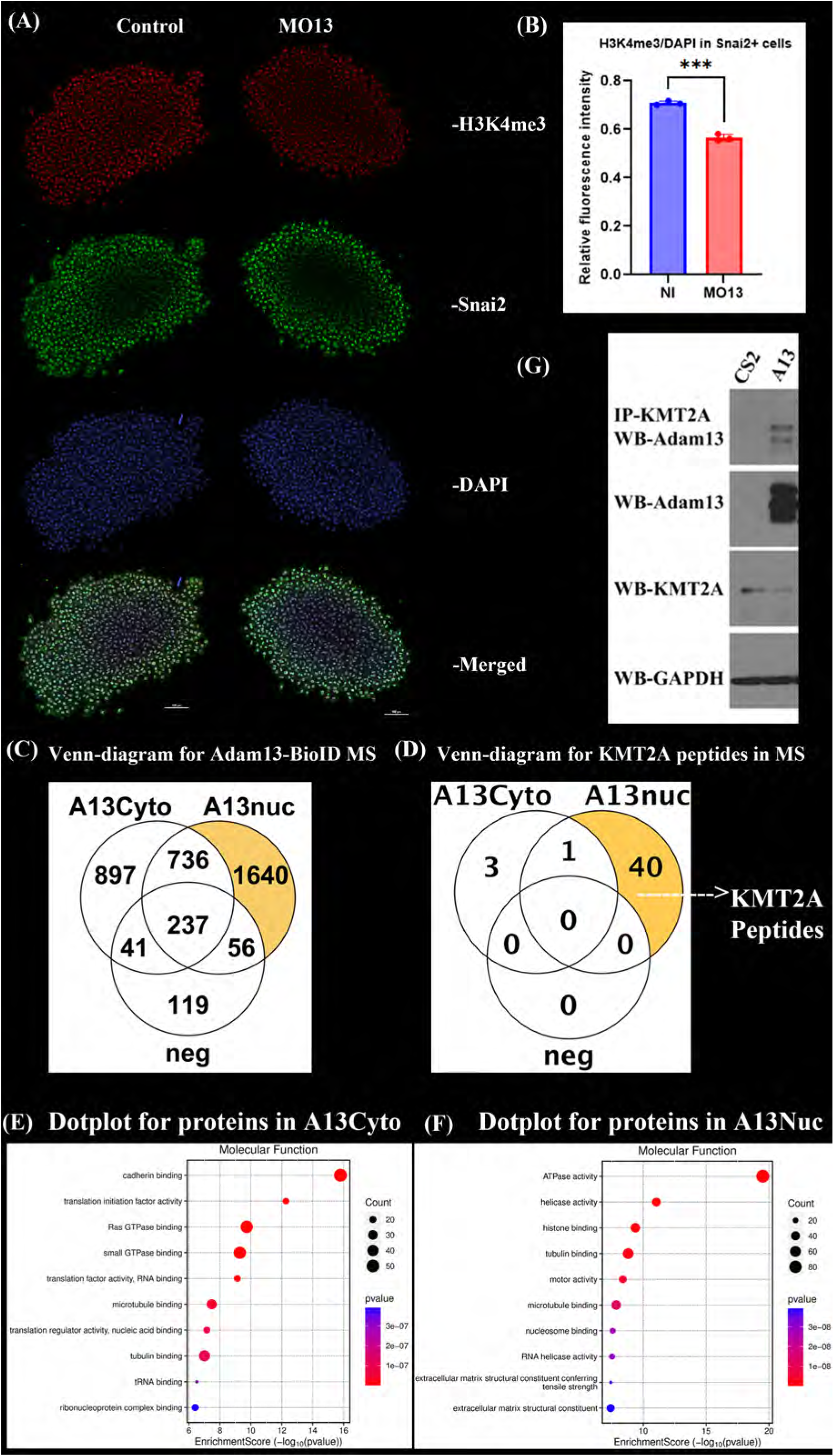
Loss of Adam13 leads to decreased H3K4me3 methylation in CNC. (A) Immunofluorescence of CNC immuno-stained for H3K4me3 (red), Snai2 (green), and DAPI (blue); all images are Max-IP. (B) Histogram depicting the relative fluorescence intensity of H3K4me3 in Snai2 positive cells between NI and MO13 CNC explants. *** p-value<0.001. (C) Venn diagram representing total proteins detected with 2 unique peptides minimum from 6 independent protein samples each for Control (neg), Adam13 cytoplasmic fraction (A13Cyto), and Adam13 nuclear fraction (A13nuc). (D) Venn diagram representing exclusive peptides detected for KMT2A in Control (neg), A13 Cytoplasmic, and Nuclear fractions. (E) Dot plot representing the gene set enrichment analysis (GSEA) for molecular function of proteins that were found exclusively biotinylated by Adam13 in the cytoplasmic fraction. The red color represents higher significance (lower p-value), and the size of the dot represents the number of genes in the set. (F) Dot plot representing the GSEA for molecular function of proteins that were found exclusively biotinylated in nuclear fraction. (G) Adam13 binding to endogenous human KMT2A was confirmed using co-immunoprecipitation in HEK293T cells. IP KMT2A, western blot Adam13, CS2 (empty vector) and Adam13 (A13, line 1). The total extract was blotted with anti-Adam13 antibody (line 2), KMT2A (line 3), and GAPDH (line 4) as a loading control.

### Adam13 interacts with proteins involved in histone modification

To understand how Adam13 could affect Histone methylation, we added a BioID tag ^41,42^ in the cytoplasmic domain of Adam13 to label proteins in close proximity (10 nm) of Adam13. We transfected HEK293T cells with either Adam13BioID or Adam13 (negative control), incubated for various times with Biotin-containing media, and isolated protein from the cytoplasm and the nucleus prior to purification with Streptavidin magnetic beads (Fig.2C). We analyzed the proteins that were significantly enriched in the cytoplasmic fraction (6 samples 897 proteins) or the nuclear fraction (6 samples 1640 proteins) using string database ^43^ to remove any doublets and pseudo proteins with no description. This led to a final list of 783 proteins in the cytoplasmic fraction and 1369 in the nuclear fraction (Fig.2C, Supplementary Fig.S2A, Supplementary table S1). This process was also followed for all mass spectrometry data in this manuscript. We analyzed the molecular function of the proteins in the cytoplasmic fraction, nuclear fraction, or common to both using Gene Set Enrichment Analysis (GSEA) pathway enrichment analysis (Fig.2E, 2F & Supplementary Fig.S2B) ^44^. The enriched molecular function for the proteins in the cytoplasmic fraction was cadherin binding (51 proteins), a known function of Adam13 and other ADAMs (Figure 2E) ^4,45^. We also found multiple SH3-containing proteins ^46,47^ including multiple Myosins and PACSIN2, the first identified ligand for Adam13 ^48^. We then looked at the enriched molecular function of proteins found in the nuclear fraction; the third most enriched function was histone binding (Fig.2F, 45 proteins). We then conducted an analysis to identify proteins associated with histone modification and found several histone demethylases (KDMs) that are known to remove methyl groups from histone H3 at lysine 9 (H3K9), including KDM3B and KDM4B, identified in the nuclear fraction (Supplementary Fig.S2C). In addition, we detected KMT2A, a protein known to methylate histone H3 at lysine 4 (H3K4), in both cytoplasmic and nuclear fractions. When we analyzed the number of exclusive unique peptides that were detected for each fraction, we found that the nuclear fraction had 40 exclusive peptides compared to 3 in the cytoplasmic fraction (Fig.2D). To confirm this binding, we performed co-immunoprecipitation using endogenous KMT2A from HEK293T cells and found it associated with transfected Xenopus Adam13 (Fig.2G). We used alphafold3 to model the potential interaction of the Adam13 cytoplasmic domain and Kmt2a (Supplementary fig.S2D). We found that all 5 models placed the interaction near the SET domain (IpTM 0.3, Fig.S2D Red). We then performed modeling of the cytoplasmic domain of Adam13 with the SET domain only (Supplementary Fig.S2E). Again, all models are consistent (IpTM 0.28). We find that the amino acids from Adam13 that are predicted to interact (Fig.S2E, yellow) with the set domain correspond to a highly conserved sequence in an overall poorly conserved amino acid sequence of the cytoplasmic domain: 790-<u>PVN</u>V<u>V</u>R<u>P</u>L<u>RP</u>-798 (Underlined Amino acids are conserved in Xenopus, Chicken, Zebrafish and Marsupial). This sequence corresponds to the beginning of the Proline rich repeats and the first SH3 binding site. On the SET domains, there are nine amino acids that are likely engaged in the interaction (Fig.S2E, Purple) including two clusters that are part of the active site and substrate binding sites (Grey, YM<u>FR</u>ID, <u>SRV</u>, YDY<u>KF</u>). In support of this model, we found multiple SET domain containing proteins labeled by ADAM13 BioID in our mass spectrometry data (Supplementary Table S1).

Taken together, these results show that loss of Adam13 affects global levels of histone methylation toward a reduced transcriptional chromatin. They also show that Adam13 interacts with multiple enzymes that are known to modify Histones.

### Loss of Adam13 affects gene expression and Tfap2α transcription start site

We have previously shown using micro-array, that loss of Adam13 affects gene expression of CNC genes such as *tfap2α* and *calpain-8* ^13^. Since the loss of Adam13 leads to changes in histone modification at a global level in CNC, we performed CNC-specific bulk RNA sequencing to discover genes impacted by the loss of Adam13 (Fig.3). We found 2575 genes that were significantly altered in the absence of Adam13, out of which 1293 were upregulated and 1282 were downregulated (Fig.3B, Supplementary table S2, and gene description of known human orthologs Supplementary table S3), including the previously identified *tfap2α* and *calpain-8. Tfap2α* has multiple transcript variants with different possible functions, as shown in human breast cancer cell lines ^49^. In *Xenopus laevis*, there are three possible transcription starts for the L gene and four for the S gene but two of these S start sites encode the same amino acid sequence (One of the extra exons is not coding), leading to three major protein isoforms for Tfap2α.L and S in CNC (Fig.3C). We quantified the relative abundance of these isoforms using transcript junctions by sashimi plot analysis (Fig.3D, Supplementary Fig.S3A & B) and exon-specific counts in both control and Adam13 morphant CNC explants. Quantification of these isoforms show that in the absence of Adam13, *tfap2α.L*-S1/*tfap2α.S*-S1 isoforms are downregulated, *tfap2α.L*-S2/*tfap2α.S*-S2 isoforms are unaffected and *tfap2α.L*-S3/*tfap2α*.*S*-S3 isoforms are upregulated (Fig.3E & Supplementary Fig.S3C). Given that *tfap2α.L* is the most abundant form during neural crest migration (Xenbase), that the sequence similarity between L and S is greater than 97% and the effect of the loss of Adam13 on L and S isoforms is the same, we focus the downstream studies on the *tfap2α.L* alloallele. When the protein sequence of Tfap2α.L-S1 is compared to Tfap2α.L-S3, we find that the alternate exon1s encode 52 amino acids that are different for the two isoforms, while the rest of the protein is identical (Fig.3G). Given that the overall level of Tfap2α mRNA is lower following Adam13 KD, we performed IF on CNC explants to detect the Tfap2α protein using an antibody that recognizes all the variants of Tfap2α.L and Tfap2α.S as well as an antibody to Sox9 to confirm CNC identity (Fig.3H). We found that Tfap2α protein levels are significantly decreased following the loss of Adam13 in neural crest cells (Fig.3F), consistent with the log fold change detected in CNC RNA-Seq (Supplementary Table S2).

**Figure 3.**
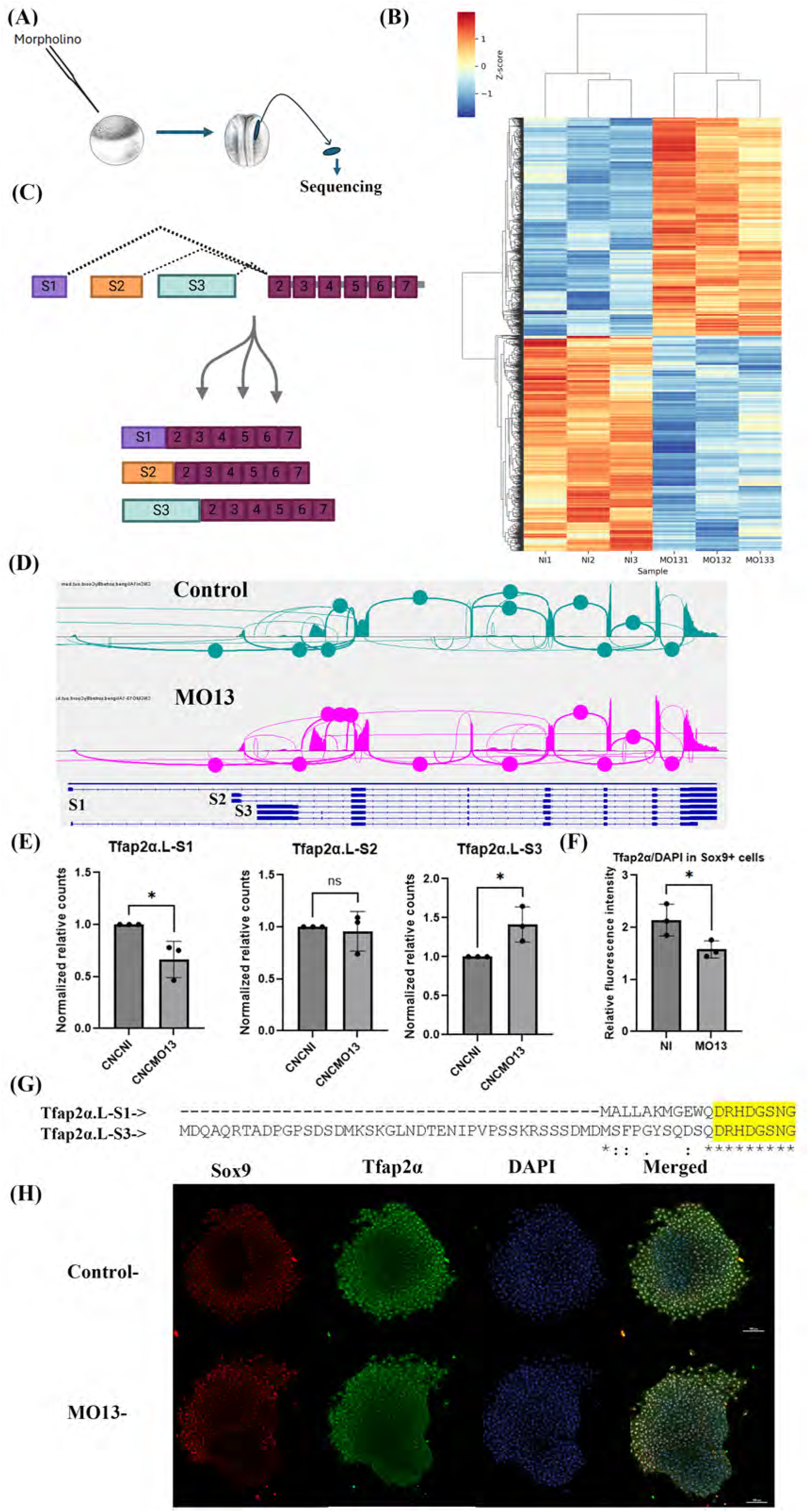
Loss of Adam13 affects gene expression and Tfap2α transcription start. (A) Schematic representation of the approach used for CNC-specific RNA-seq. Control (NI) and Adam13KD (MO13) CNC were dissected at the same apparent stage (Stage 17) in 3 independent experiments. (B) Heatmap visualization of RNA-sequencing data between Controls (NI1, NI2, NI3) and Adam13KD (MO131, MO132, MO133), higher Z-score in red representing increase expression and blue representing decreased expression. (C) Schematic representation of the Tfap2α.L gene. Three putative transcription start sites (alternate exon1) connecting to exon2 and producing three different proteins differing at the N-terminus. (D) Sashimi plot analysis of transcript junctions detected for the Tfap2α.L gene in RNA-sequencing data between Control (NI1) and Adam13KD (MO131), circles mark the major transcript junction between corresponding exons. (E) Quantification of exon-specific counts for the three Tfap2α.L alternate starts (S1, S2 and S3) normalized to the total transcript counts for the Tfap2α.L gene. Student t-test was performed for statistical analysis, *=p-value<0.05; ns=p-value>0.05. (F) Histogram depicting relative fluorescence intensity of Tfap2α in Sox9 positive cells between Control (NI) and Adam13KD (MO13) CNC explants. *=p-value<0.05. (G) ClustalW protein sequence alignment for Tfap2α-S1 and S3 to highlight the sequence difference in exon1 and the yellow highlighted sequence is Exon2 which is common. (H) Immunofluorescence of CNC explants immuno-stained for Sox9 (red), Tfap2α (green) and DAPI (blue) and imaged using confocal microscopy, all images are Max-IP.

Taken together, these results show that the loss of Adam13 in the CNC affects multiple gene expression levels. They further show that both Tfap2α mRNA and protein levels are reduced, and that the transcription start is shifted in the absence of Adam13 to a different exon.

### Adam13 controls Arid3a binding and histone methylation at the Tfap2α.L promoter

To generate large numbers of embryos missing the Adam13.L protein, we used CRISPR-Cas9 to generate *Xenopus laevis* knockout (KO, NXR, MBL). Homozygotes adult frogs are viable, similar to what is seen for most ADAM KO performed in mice ^50–52^. We used embryos generated from either Wild Type (WT) or KO frogs to test whether Adam13 affected the epigenetic landscape on the *tfap2α* gene promoter. We performed Chromatin Immuno-Precipitation (ChIP) at stage 20, followed by qPCR using primers against the promoter region of *tfap2α.L*-S1 (S1) and *tfap2α.L*-S3 (S3). For simplification we will refer to the 2 promoter and splicing variant as S1 and S3 in the rest of the text. We first tested the relative abundance of DNA associated with H3K4me3 and found that Adam13 KO leads to decreased DNA associated with H3K4me3 for the S1 promoter and an increase for the S3 promoter (Fig.4A & Supplementary Fig.S4A). We next performed ChIP-qPCR for H3K9me2/3, and we found that there is an increase of DNA associated with H3K9me2/3 at the S1 promoter but no change at the S3 promoter (Fig.4B & Supplementary Fig.S4B). Given that Adam13 and Arid3a regulate the expression of *tfap2α* ^6^, we tested whether Arid3a binding to the *tfap2α* promoter depends on the presence of Adam13. In the absence of Adam13, there was less DNA associated with Arid3a for the S1 promoter, while the loss of Adam13 had no significant effect on DNA associated with Arid3a for the S3 promoter (Fig.4C & Supplementary Fig.S4C). Finally, we tested if Adam13 was present as part of the complex at the promoter region by performing ChIP using a cytoplasmic domain-specific antibody for Adam13 ^13^. We find that Adam13 is present at both the S1 and S3 promoters (Fig.4D & Supplementary Fig.S4D).

**Figure 4.**
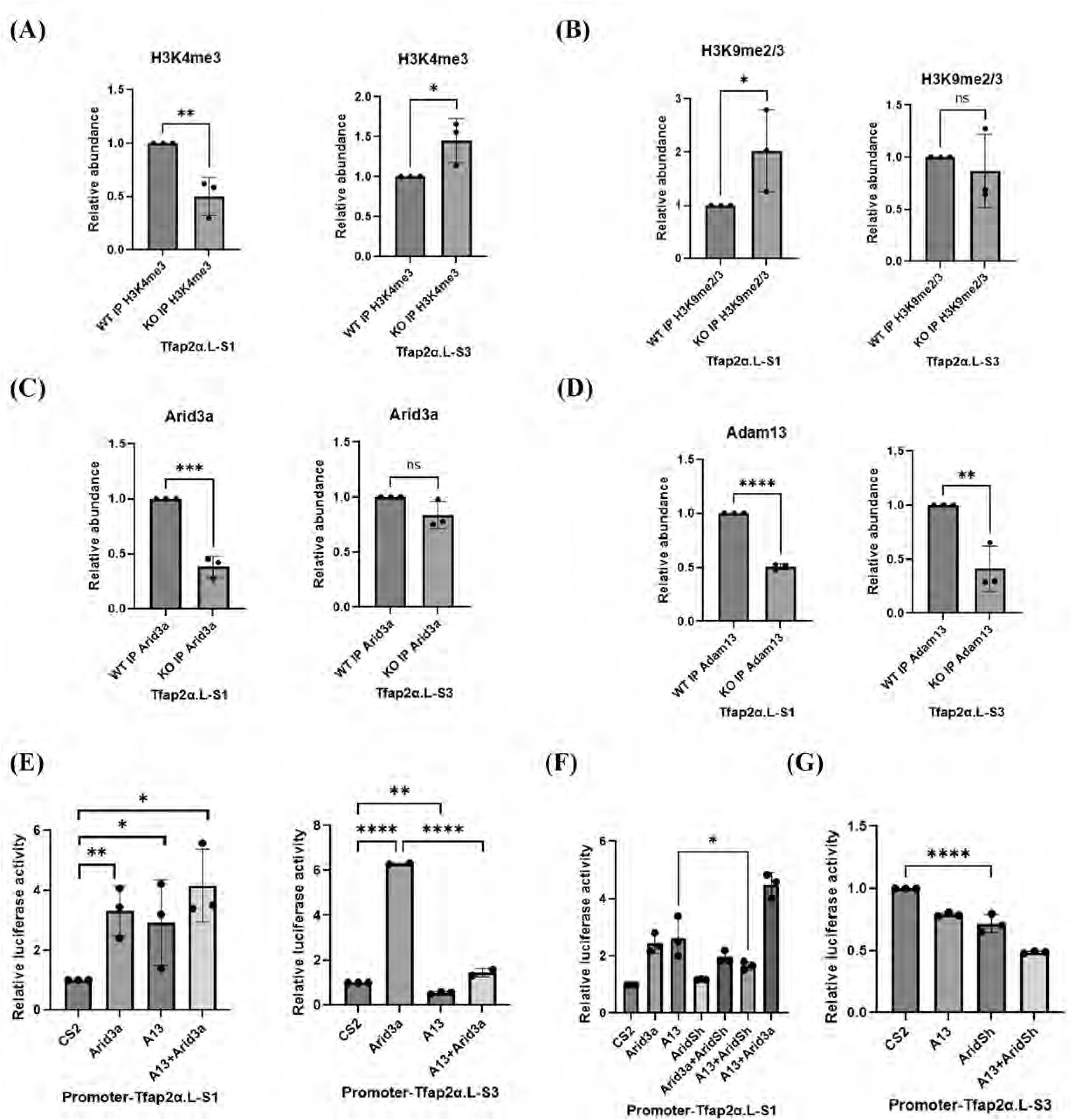
Adam13 controls Arid3a binding and histone methylation at the Tfap2α.L promoter. Chromatin-Immunoprecipitation-qPCR analysis of DNA associated with the chromatin binding proteins. (A, B, C, D) Histogram depicting the relative abundance of DNA bound to H3K4me3 (A), H3K9me2/3 (B), Arid3a (C), and Adam13 (D) in WT and Adam13KO embryos at the promoter region of Tfap2α-S1 and S3 based on ChIP-qPCR analysis. ChIP was performed at stage 20. Student t-test was performed for statistical analysis, * p-value<0.05, ** p-value<0.01, *** p-value<0.001, **** p-value<0.0001, ns represents p-value>0.05. (E-G) Histogram depicting the relative luciferase expression using the luciferase reporter containing Tfap2α-S1 or S3 promoter region in HEK293T cells that were transfected with CS2 (empty vector), Arid3a, Adam13, Arid3a+Adam13, alone or in combination with the Human Arid3a-SH (AridSh, F-G), along with the CMV renilla reporter. One-way ANOVA was performed for statistical analysis. * p-value<0.05, ** p-value<0.01, **** p-value<0.0001.

The results obtained for the S1 promoter with Histone modification reflect the overall changes observed in the CNC (Fig.1&2), namely that the loss of Adam13 increases repressive marks (H3K9me2/3) and decreases activating marks (H3K4me3). In addition, these results show that Adam13 localizes to the promoter of both transcription starts but only regulates Arid3a association to the S1 start site.

To test the impact of Adam13 and Arid3a on the *tfap2α* promoter, we cloned the upstream region of *tfap2α.L*-S1 (S1, 2368 bp) and *tfap2α.L*-S3 (S3, 3577 bp) with a luciferase reporter gene and tested the luciferase reporter expression in HEK293T cells (Fig.4E). We find that transfection of Arid3a and Adam13 increase the luciferase expression alone and that the expression of both proteins further increases the level of luciferase activity for the S1 reporter. However, the S3 reporter increased in the presence of Arid3a but significantly decreased in the presence of Adam13 (Fig.4E). This is reminiscent of the observation obtained in the CNC RNAseq data showing that loss of Adam13 reduces S1 usage while increasing S3 usage (Fig.3E).

We next tested whether the increase in the luciferase expression by Adam13 alone for S1 was due to endogenous *ARID3A* using a shRNA against human *ARID3A* co-transfected with Adam13. We found that KD of endogenous *ARID3A* expression significantly decreased S1 reporter luciferase expression by Adam13 (Fig.4F). We also saw a significant decrease in the S3 reporter luciferase expression when endogenous *ARID3A* was knocked down (Fig.4G).

Taken together these results show that Adam13 localizes to the S1 promoter and regulates both Arid3a binding and Histone modifications. Adam13 binding to the S1 promoter induces increased transcription in an Arid3a dependent manner. In contrast, Adam13 binding to the S3 promoter reduces transcription, reduces H3K4me3, and has no significant effect on Arid3a binding or H3K9 methylation status.

### Tfap2α variants have distinct biological functions

We have previously shown that the loss of Adam13 affects CNC cell migration *in-vivo* ^3,53^; we then tested whether we could rescue CNC cell migration in Adam13 KD embryos using mRNA encoding the Tfap2α.L-S1 or S3 variants. To do this, we used an 8-cell targeted injection of mRNA combined with *adam13* morpholino (MO13) in the dorsal animal blastomere of the Xenopus embryo to target prospective neural crest cells (Fig.5A). This method allows for a rapid binary count assessing for the presence or absence of fluorescent CNC in the migration pathway. Each injection is calibrated by injecting the lineage tracer alone to calculate targeting efficiency, which varies from female to female due to the quality of pigmentation (50 to 90%). We have shown that this method faithfully reproduces the results obtained using CNC graft, a much more labor-intensive technique ^4,8,13,53,54^. We used Membrane Cherry (MbC) as our lineage tracer. When co-injected with the *adam13* morpholino (MO13), migration was observed in only 53% of the embryos. This decrease in cell migration was rescued by the *tfap2α.L*-S1 mRNA in 81% of the embryos, while *tfap2α.L*-S3 mRNA was only able to rescue up to 66% of the embryos (Fig.5B) using the same conditions and dose of the mRNAs.

**Figure 5.**
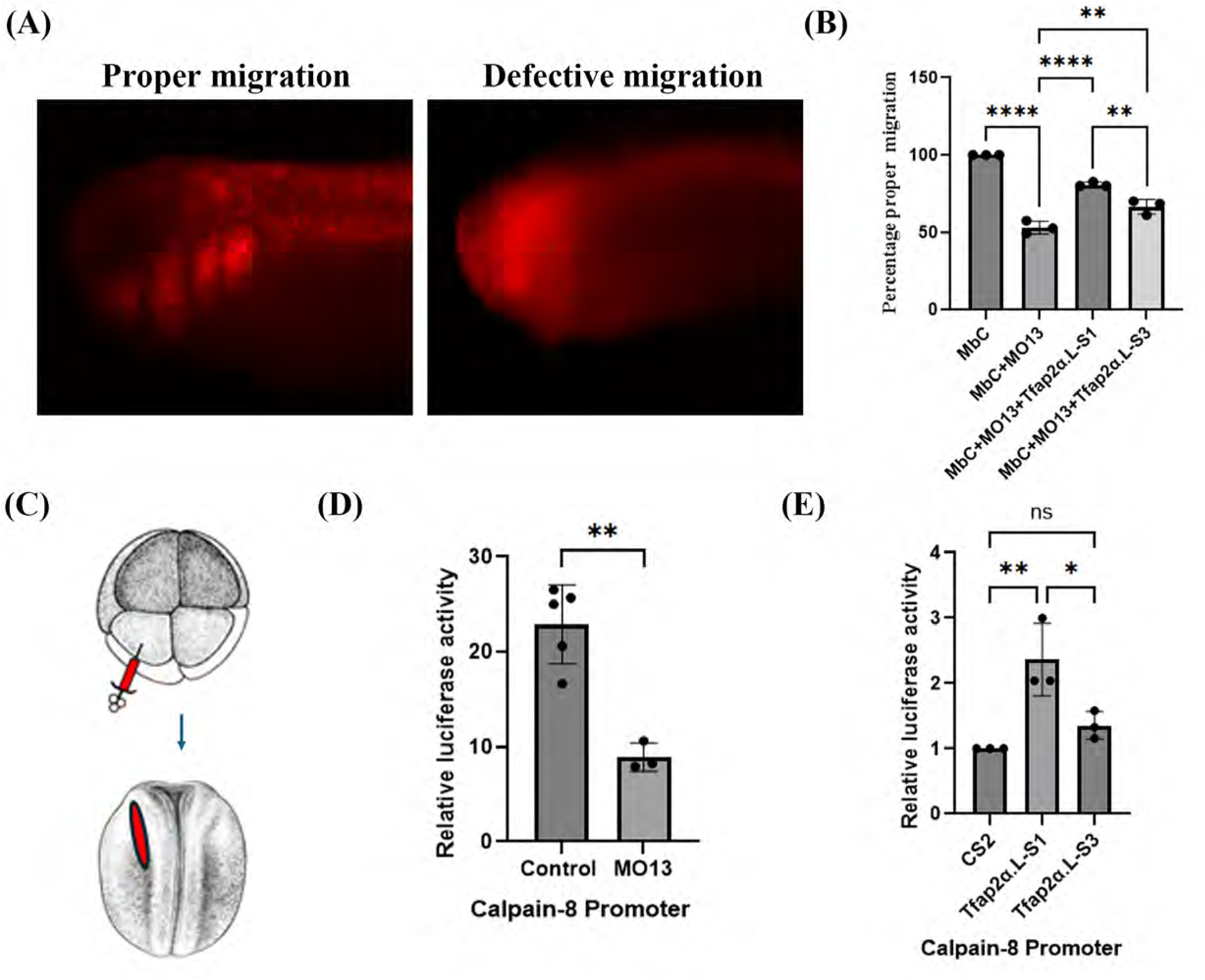
Tfap2α variants have distinct biological functions. (A) Fluorescence tracking of CNC cell migration. Embryos were injected with fluorescent Membrane Cherry (MbC) mRNA at the 8-cell stage in one dorsal animal blastomere and scored at stages 22-24 for the presence or absence of fluorescence in the branchial arches to quantify migration. (B) Histogram representing the percentage proper migration in Control (MbC), Adam13KD (MO13), MO13+Tfap2α-S1, MO13+Tfap2α-S3. One-way ANOVA was performed for statistical analysis. ** p-value<0.01, **** p-value<0.0001. (C) Schematic representation of the strategy for luciferase assay in embryos. Embryos were injected at the 8-cell stage in the dorsal blastomere with Calpain-8 luciferase reporter and the CMV renilla reporter to target prospective CNC. Embryos with an expression of MbC in the CNC region at stage 18 were selected to perform luciferase assay. Individual embryos were used for luciferase reading. (D) Histogram depicting relative *calpain-8* luciferase reporter expression normalized to CMV renilla expression in Control and Adam13KD (MO13) embryos. Student’s t-test was performed for statistical analysis. ** p-value<0.01. Each experiment was repeated 3 times with 5 individual embryos per case in each experiment. (E) Histogram depicting the relative *calpain-8* luciferase reporter expression in cells transfected with CS2 (empty vector), Tfap2α.L-S1-Flag, and S3-Flag along with Calpain-8 luciferase reporter and the CMV renilla. One-way ANOVA was performed for statistical analysis. * p-value<0.05, ** p-value<0.01, ns p-value>0.05.

Given our previous results showing that Adam13 regulates *calpain-8* through Tfap2α ^6^, we used a luciferase reporter with the Xenopus *calpain-8* promoter (2.5 kbp) to test the activity of the Tfap2α variants. We first assess whether this reporter could respond to Adam13 *in vivo* in the CNC by injecting the reporter with either MbC or MbC and MO13 at the 8-cell stage in the CNC precursor. We selected embryos at stage 20 that expressed the fluorescent marker in the CNC to perform luciferase reading (Fig.5C). We saw a significant decrease in *calpain-8* luciferase expression in the absence of Adam13 (Fig.5D) aligned with what was observed for the endogenous *calpain-8* gene expression ^13^, confirming that the promoter responds to Adam13. To test whether both Tfap2α isoforms could activate *calpain-8*, we transfected the *calpain-8* luciferase reporter with CS2 (empty vector), Tfap2α.L-S1-Flag or Tfap2α.L-S3-Flag. We found that Tfap2α.L-S1-Flag significantly increased the luciferase expression while Tfap2α.L-S3-Flag did not (Fig.5E).

### Tfap2α.L-S1 and Adam13 regulate splicing

To test if proteins associated differently with the two Tfap2α isoforms, we inserted a Flag tag at the c-terminus of Tfap2α.L-S1 and S3) and used an anti-flag antibody to IP both proteins from HEK293T cells to performed mass spectrometry. HEK293T cells transfected with RFP-Flag served as a negative control to eliminate any proteins binding non-specifically to either the beads or the anti-Flag-antibody. We identified 65 proteins that were significantly associated with Tfap2α-S1-Flag but not S3, and 28 proteins with Tfap2α-S3-Flag, while 1772 were not significant (Fig. 6A, Supplementary table S4). When looking at proteins that were present in both samples with a minimum of two unique peptides, we found 506 proteins. We next performed GSEA for the biological process of the 65 identified proteins and found RNA splicing to be the most enriched biological process (Fig.6B, 44 proteins). While Spicing and RNA binding were also found enriched in the list of common proteins, this pathway was not enriched in the protein uniquely associated with the S3 isoform.

**Figure 6.**
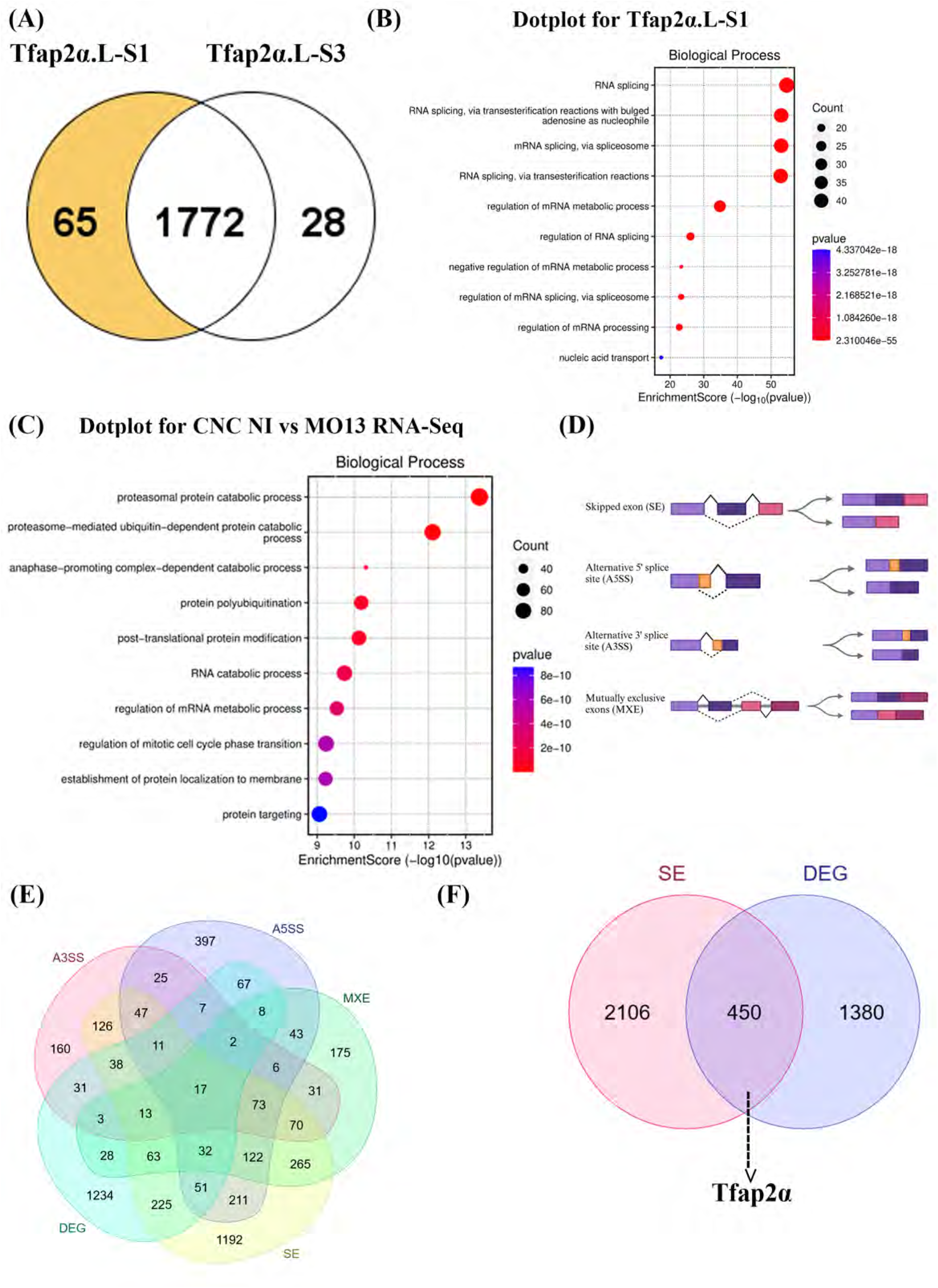
Tfap2α.L-S1 and Adam13 are key regulators of splicing. (A) Venn diagram representing proteins detected following IP of Tfap2α.L-S1-Flag and Tfap2α.L-S3-Flag after removing proteins found in the RFP-Flag negative control. We used T-test to identify proteins that were significantly enriched in either of the 2 samples. Proteins that associate significantly more with the S1 variant (65) are highlighted in yellow. (B) Dot plot representing the gene set enrichment pathway analysis for biological processes of proteins that were found significantly associated with Tfap2α-S1-Flag. The red color represents higher significance (lower p-value), and the size of the dot represents the number of genes identified in the set. (C) Dot plot representing the gene set enrichment pathway analysis for biological processes of differentially expressed genes in Adam13KD (MO13) CNC. (D) Schematic representation of the different types of alternative splicing events detected by rMATS analysis on CNC RNA sequencing data. (E) Venn diagram depicting the genes found to have significant differences in splicing in CNC lacking Adam13. Genes having Alternate 5’ splice site (A5SS), Alternate 3’ splice site (A3SS), and mutually exclusive exons (MXE) are compared to differentially expressed genes (DEG). (F) Venn diagram depicting the genes found statistically differentially expressed versus genes having skipped exon (SE).

Given that the S1 isoform expression depends on Adam13, we re-analyzed the CNC RNAseq data to test if overall RNA splicing was affected by the absence of Adam13. We looked at biological processes enriched for the differentially expressed genes and found RNA catabolic process (78 proteins) and regulation of mRNA metabolic process (68 proteins) among the top 10 enriched biological processes (Fig.6C). We also identified all the genes related to RNA splicing and represented them in a custom heatmap (Supplementary Fig.S5A).

To distinguish between the differentially expressed and differentially spliced genes, we used rMATS ^55^, which quantifies the different possible splicing events in the RNA-sequencing data, as shown in (Fig.6D). We quantify the differentially spliced events including alternative 3’ splice sites (A3SS), alternative 5’ splice sites (A5SS), skipped exons (SE), and mutually exclusive exons (MXE), and selected the genes that were statistically different between the control and Adam13KD CNC (PSI value difference: |ΔPSI| ≥0.05, False discovery rate: FDR ≤0.01). The list of genes that were differentially spliced was then compared to the list of genes that were differentially expressed (Supplementary table S5). We compared genes in all different splicing events to the list of differentially expressed genes and represented them in Venn diagrams (Figure 6E, F & Supplementary Figure S5B).

This analysis shows that there is a partial overlap (596) between genes that are differentially expressed (1,830 genes) and those that are differentially spliced (3539) in CNC lacking Adam13 (Supplementary Fig.S5C). Among the splicing events, exon skipping was the most affected in the absence of Adam13 (2,556 genes). *Tfap2α* is one example of a gene that is both alternatively spliced (Exon-skipping) and differentially expressed in a list of 450 genes. Taken together, our results show that Adam13 positive regulation of Tfap2α-S1 variant is critical for CNC migration, expression of *calpain-8,* and RNA splicing in the CNC.

### ADAM interact with RNA binding proteins

To test if our results with Adam13 could be generalized to other ADAM we performed IP from HEK293T cells using an antibody to endogenous human ADAM9 using normal Rabbit immunoglobulin as a negative control. We selected ADAM9 given its partially overlapping function with Adam13 to promote CNC migration ^4^, and its known function in the nucleus ^16^. As expected, we found between 5 and 7 unique peptides per sample corresponding to ADAM9 confirming the specificity of the antibody. We found 166 proteins that were significantly associated with ADAM9. When we compared the proteins found in the ADAM9 IP to the proteins found in Adam13 IP and Adam13BioID we obtained a list of 57 proteins (Supplementary table S6). We performed string analysis with the 57 proteins (44 were identified in String including ADAM9 and ADAM13/33) and found that the most enriched biological processes were RNA processing (Fig.7A red) and mRNA splicing via the spliceosome (Fig.7A blue). The molecular function analysis shows RNA binding (Fig.7A green). Analysis using GSEA confirms these finding (Fig.7B). Given the number of RNA binding proteins in this list we hypothesized that IP of Adam13 and ADAM9 may contain RNA if the complexes are stable enough. To test this, we extracted transfected and non-transfected HEK293T cells in the presence of RNAse inhibitor and performed the same IP as for the Mass spectrometry. Our results show that Adam13 but not the mutant lacking its cytoplasmic domain co-precipitate with RNAs (Fig.7C). Similarly, we found that endogenous ADAM9 also precipitated RNAs (Fig.7C). Given that the proteins labeled by Adam13BioID also contained multiple RNA-binding proteins, we tested if they could bring down RNAs as well. Indeed, when protein extracts from cells transfected with Adam13BioID or Adam13 were purified using streptavidin-magnetic beads, RNA was found exclusively in the Adam13BioID samples (Fig.7C). To test if we could also co-precipitate Adam13 in embryos with RNA, we used wild type and Adam13KO embryos and immunoprecipitated with a monoclonal antibody to the extracellular domain (mAb6A4) ^56^. The result show that Endogenous Adam13 co-precipitate with RNA in Xenopus embryos (Fig.7D).

**Figure 7.**
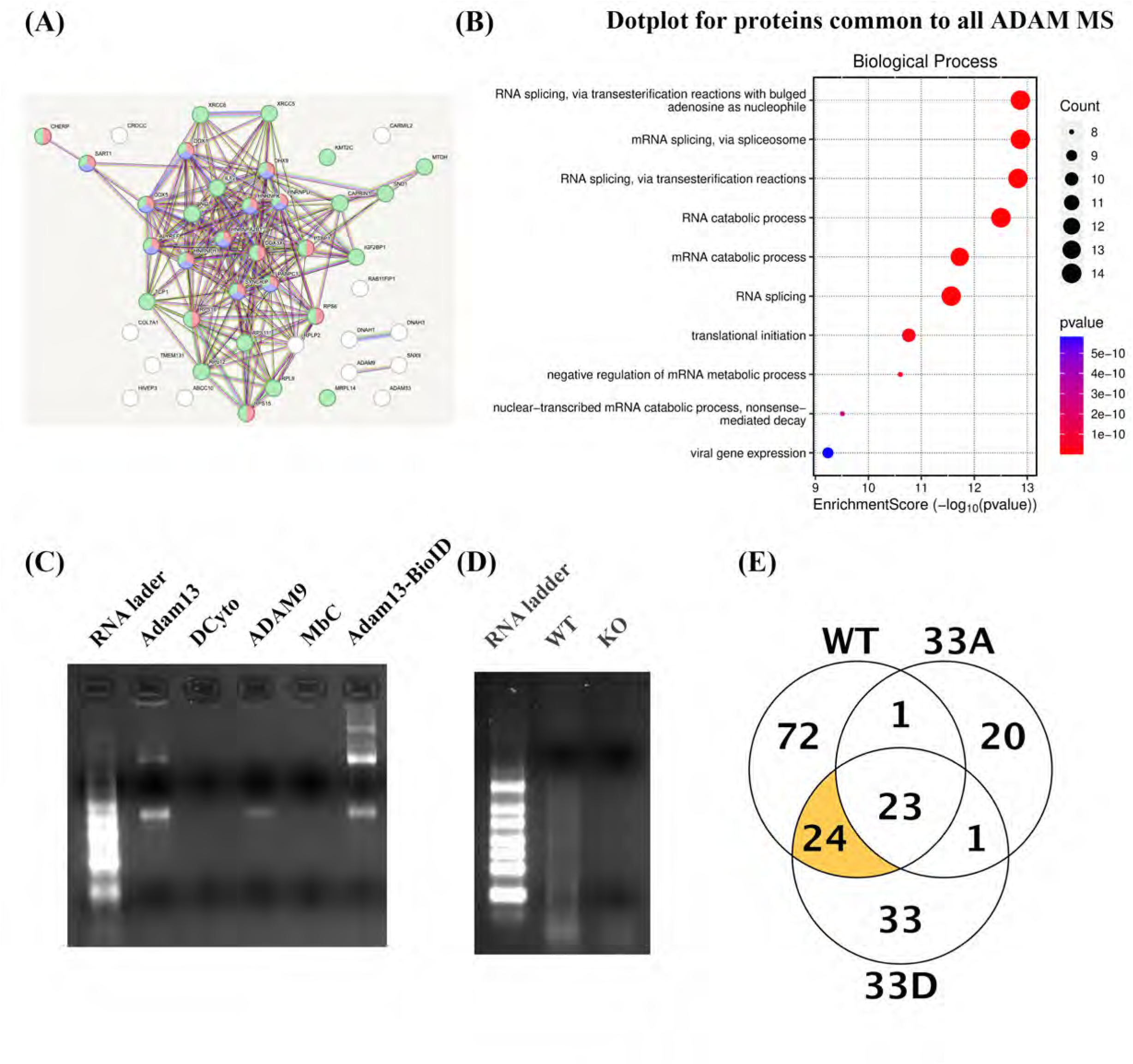
ADAM proteins interact with RNA binding proteins and their RNA cargo. (A) String plot of proteins identified in ADAM9, Adam13 IP and Adam13-BioID. Red indicates proteins involved in RNA transport, blue mRNA splicing via spliceosome (Biological processes G.O.) and Green RNA binding (Molecular function G.O.). Lines between protein indicate previously identified links. (B) Dot plot representing the gene set enrichment pathway analysis for biological processes of proteins that were found significantly associated with ADAM9 and 13. (C-D) Agarose gel of RNA purified by Immunoprecipitation. Lane 1 RNA marker, Lane 2 and 3 RNA immunoprecipitation using a monoclonal antibody to the Adam13 extracellular domain from HEK293T cells transfected with Adam13 (2) or the mutant lacking the cytoplasmic domain (DCyto, 3). RNAIP from non-transfected HKE293T cells with an antibody to human ADAM9 (Lane 4). Lanes 5 and 6 correspond to Streptavidin pull down of proteins from cells transfected with Membrane Cherry (MbC) or Adam13-BioID. (D) RNA IP using an Adam13 monoclonal antibody from 10 wild type (WT) embryos or Adam13 KO (KO) embryos.

We have previously shown that Adam13 is phosphorylated by GSK3 and polo like kinase and that this last phosphorylation at Threonine 833 is critical for Adam13 function in the CNC ^54^. Phosphorylation by Polo like kinase does not affect protein stability, proteolytic activity or subcellular localization. We used Mass spectrometry to identify proteins that could bind to Adam13, the phosphomimic Adam13-833D but not the non phosphorylatable mutant Adam13-833A. We found 23 proteins that were common to all 3 proteins, and 24 that were unique to Adam13 and Adam13-833D (Fig.7E, Supplementary table S7). While the 23 protein that were common to all included multiple RNA binding proteins, only the proteins exclusive to Adam13 and Adam13-833D contained protein involved in mRNA transport and Splicing via the spliceosome (SRSF1, 3 and 7, DHX9 and HNRNPA2B1) reinforcing the notion that this aspect of Adam13 activity is critical for CNC migration.

Together these results show that Adam13 and ADAM9 bind to RNA-binding proteins and their RNA cargo. For Adam13 this depends on the presence of its cytoplasmic domain. This is not unique to Adam13 or due to overexpression as endogenous ADAM9 protein also associate with RNA containing complexes. In Addition, phosphorylation by Polo like kinase promotes binding to proteins involved in mRNA transport and splicing.

## Discussion

The role of ADAM proteins in neural crest cell migration is well documented in amphibian, chick, and, to a lesser extent in mice ^45,57^. The lack of obvious results in the mouse can be attributed partly to the overlapping functions of ADAM proteins and their widespread expression, which leads to multiple organs phenotypes. Our group has shown that Adam9, 13 and 19 cooperate to promote cranial neural crest cell migration ^5,13,53^. The proteolytic activity of Adam9 and 13 can cleave the same substrate such as Cadherin-11 and protocadherins. We have shown that the cleavage of these cell adhesion molecules produces extracellular fragments that are pro-migratory ^4,6,7^. Similar activities were identified for Adam proteins in the avian embryo where Adam10 and Adam19 cleavage of multiple cadherins is essential for the neural crest EMT ^9,45^. In mice, Adam19 was found to be critical for the proper positioning of Cardiac neural crest to form the heart ^58^. In most species, Adam10 is critical for neural induction given its role as notch alpha-secretase and is also likely to contribute to neural crest specification ^59^. This is difficult to show given the wide range of embryonic defects in mice lacking Adam10 ^11,60,61^. In Xenopus, Adam13 was found to cleave ephrin resulting in a local activation of the Wnt signaling pathway important for neural crest induction ^62^. In Xenopus and Chick, Adam13 is expressed in cells that migrate ahead of the flow (pioneer cells) suggesting a role in either opening the pathway or detecting/modifying cues to help cells follow the same path ^53,63^. We have shown that in Xenopus, Adam13 is critical in the leading-edge cells and that the physical opening of the space between the mesoderm and the neural crest is sufficient to rescue the loss of Adam13 ^53^. Non-proteolytic ADAM proteins are also involved in CNC specification and migration. In particular, Adam11 binds to BMP4 and its receptor increasing the local response and regulating cell proliferation and neural tube closure. In addition, loss of Adam11 results in an increase in Wnt signaling characterized by increased nuclear localization of β-catenin and precocious EMT leading to abnormal CNC migration ^2^.

### Role of ADAM proteins in the nucleus

Three ADAM proteins have been found in the nucleus, ADAM9, 10, and 13 ^13,16,64^. While the role of ADAM10 in the nucleus has not been studied extensively, a recent study of ADAM9 in cancer cells demonstrated that the protein is associated with the promoter of genes that typically inhibit angiogenesis. The study also showed that ADAM9 functions as a repressor on these promoters, resulting in the expression of pro-angiogenesis genes ^16^. However, the mechanism by which ADAM9 regulates gene expression remains unclear. In Xenopus, we have shown that the Adam13 cytoplasmic domain is cleaved by gamma-secretase and translocates into the nucleus ^13^. On its own, the cytoplasmic domain does not activate gene expression ^6^. We have also shown that Adam13 interacts with the transcription factor Arid3a, inducing a proteolytic cleavage and resulting in the activation of multiple genes including the key transcription factor Tfpa2α ^6^. Loss of Adam13 in the CNC results in modest changes in the expression of a large number of genes ^13^. In this report, we have shown that loss of Adam13 decreases the overall level of H3K4 trimethylation (Fig.2) while increasing the H3K9 di-trimethylation (Fig.1) suggesting that Adam13 may promote a more transcriptionally active chromatin. Mass spectrometry results show several SET-domain containing proteins associated with Adam13 including enzymes known to add methylation on the lysine 4 of histone H3 (KMT2A, Fig.2) and remove methylation on lysine 9 of histone H3 (KDMs, Fig.2). Focusing on the Tfpa2α promoter, we find that the loss of Adam13 reduces H3K4 and increases H3K9 to the promoter of the most upstream ATG (Fig.4). These changes in methylation are associated with a decreased binding of Arid3a to this site and the decreased transcription of Tfpa2α at this position, suggesting a clear picture where a complex composed of Adam13, Arid3a and the KMT and KDM enzymes bind to the promoter of selected genes to increase their transcription and/or produce alternate isoforms. The luciferase reporter experiments suggest that Adam13 and Arid3a can regulate the transcription of a plasmid (Fig. 4), suggesting that they can function independently of Histone modification when open DNA is accessible. This suggests that Histone modification could happen first, allowing the Adam13/Arid3a complex to subsequently regulate transcription of the open chromatin. The results corresponding to the third transcription start of Tfpa2α show that Adam13 can act both as an activator and a repressor depending on the context, in line with our observation that a similar number of genes are significantly up or downregulated in the CNC lacking Adam13 (Fig.3B). The fact that endogenous human ADAM9 also interacts with multiple KMT2 suggest that post translational Histone modification is a general function of ADAM proteins and may contribute to its identified repressive role in human cancer ^16^.

We have found that Adam13 regulates splicing in the CNC. The number of genes that are differentially spliced (2,556) in the absence of Adam13 is larger than the number of genes that are simply differentially expressed (1,830). Based on our findings, Adam13 could affect splicing at multiple levels. First, as shown for Tfap2α, change in histone methylation at alternate transcriptional start leads to different exon usage, in fact, there is increasing evidence that local histone modifications can affect splicing ^65^. In-particular as we have shown for Tfap2α, presence of H3K9me3 can lead to changes in the inclusion of alternative exons ^66^. Second, Adam13 induction of the Tfap2α-S1 transcript, which seem to be associated with unique proteins involved in Splicing (Supplementary Table S4), could also be responsible for the specific splicing pattern present in the CNC (Discussed more below). Finally, Adam13 protein could be directly involved in splicing given its association with proteins that are part of the spliceosome complex. The fact that the form of Adam13 that is phosphorylated by Polo like kinase and is critical for CNC migration ^54^ appears to uniquely associate with proteins involved in splicing suggest that this function is essential and is tightly regulated (Supplementary Table S7). Our Mass spectrometry results of human ADAM9 shows association with KMT2A, while among the proteins that bind to both ADAM9 and Adam13 are proteins involved in RNA splicing (PABPC1, RNA helicases, PTBP1, HNRNPs, SART1) as well as histone modification (KMT2C). This suggest that ADAM proteins in general may be found in multiprotein complexes associated with RNA to participate in RNA trafficking, splicing or regulation of translation. Additional studies are needed to address this possibility.

### Biological relevance of the Tfap2α isoforms

The presence of Adam13 at the Tfap2α promoter results in the expression of one isoform of Tfap2α from the most upstream start site (S1) and the repression of another isoform at the third potential start site (S3). The two proteins differ in the first 52 amino acid at the N-terminus of the proteins. While these 52 amino acids do not encode an obvious critical domain, the biological activity of these two isoforms is clearly different with the S1 isoform able to rescue CNC migration in the absence of Adam13 much more efficiently than the S3 isoform, most likely due to the ability of S1 to induce the *calpain-8* promoter while S3 does not (Fig.5). This suggests that the S1 isoform is uniquely adapted at regulating gene expression critical for the CNC. Our Mass spectrometry analysis reveals that several proteins involved in RNA splicing associate significantly more with the S1 isoform (Fig.6). This observation could explain how the loss of Adam13 significantly affects gene splicing in the CNC (Fig.6). Tfap2α variants have been identified in humans as well. Most notably, two isoforms of Tfap2α differing in their N-terminus in a very similar manner as what we found here are expressed in Human breast cancer. In these cells, while both isoforms regulate Erbb2 gene expression, only one of the isoforms can act as a transcriptional repressor for Cyclin D3 due to one SUMOylating site present at the N-terminus ^49^. While the S1 form does not have this conserved sequence we found that the S3 isoform contains a SUMOylating site within its N-terminus (15-SDM<u>K</u>S-19). Additional experiments would be required to fully understand the functional differences of the two major isoforms of Tfap2α in Xenopus.

Adam13 appears to be part of large protein complexes involved in multiple functions that are critical for cell adhesion, cell migration, gene expression and chromatin remodeling. As unexpected as this is, we and other have found strong evidence of these different function *in vivo* in multiple animal models. Adam13 does not turn on or off genes but rather modulate the expression or splicing of existing genes ^13^. As such we propose that Adam13 could be a sensor present at the cell surface able to regulate gene expression to better adapt to changes in the extracellular environment. The number of Calcium binding proteins and ion channels that are found associated with Adam13, as well as Adam13’s ability to interact with proteins of the extracellular matrix ^12^ suggest that it could “sense” ionic variations or other ECM associated signals to trigger changes in gene expression.

## Acknowledgment

This work was supported by grant from the NIH NIDCR to D.A. R01DE016289 and office of directorate R24OD21485. M.H. is supported by P40OD010997 and R24OD030008. We thank Genevieve Abbruzzese for the Calpain-8 luciferase construct and Daniel Splittstoesser for his contribution to the project. We thank Drs. Jesse Mager and Kimberly Tremblay for confocal microscopy help. Mass spectral data were obtained at the University of Massachusetts Mass Spectrometry Core Facility, RRID:SCR_019063 (Director Stephen Eyles). The Orbitrap fusion was purchased with NIH support: Award Number S10OD010645. The content is solely the responsibility of the authors and does not necessarily represent the official views of the National Institutes of Health.

## Author contributions

DA and AP contributed to the experimental design, perform experiments and analysis and wrote the manuscript. HC contributed to embryo injection CNC dissections and phenotypical analysis. SK designed and cloned all Arid3a construct, the S3-luciferase reporter and perform targeted injection and luciferase assays. LT produced Adam13-BioID construct and perform the experiments of proximity labelling. He also contributed to bio informatic analysis. AP and AC contributed to the bioinformatic analysis. KC, NS and MH produced the Adam13KO frogs, characterized the mutation and provided adult animals used in this manuscript.

## Materiel and Methods

### Antibodies

The following antibodies were used in this study: monoclonal antibody to Ribophorin-1 (Rpn1) was used as loading control, RRID:AB_2687673. Mouse monoclonal antibody to H3K9me2/3 (Cell Signaling Technology Cat# 5327, RRID:AB_10695295). Mouse monoclonal to Arid3a (DSHB Cat# PCRP-ARID3A-1E9, RRID:AB_2618410). Rabbit monoclonal to H3K4me3 (Cell Signaling Technology Cat# 9751, RRID:AB_2616028). Rabbit monoclonal to KMT2α (Cell Signaling Technology Cat# 14197, RRID:AB_2688010). Mouse monoclonal to TFAP2α (DSHB

Cat# 3B5, RRID:AB_2202275). Mouse monoclonal to anti-flag tag (DSHB Cat# 12C6c, RRID:AB_2890618). Anti Adam13 against cytoplasmic domain and extracellular domains have been described elsewhere ^13,56^.

### Morpholinos and DNA constructs

Morpholino antisense oligonucleotides (Gene Tools, Philomath OR) were designed to block the translation of Adam13 (MO13-GTCCCAGCCGACCCTCC). *adam13.l* was cloned in pCS2 vector. The BioID2 tag ^67^ was added to *adam13.l* using Takara infusion cloning at amino acid 893 within the cytoplasmic domain (NSAT-BioID2-QLKG). All Adam13 construct were done on the “Rescue” construct that has silent mutations on the morpholino binding site. BioID construct were done on the Wild type Adam13 as well as the E/A proteolytically inactive mutant (A13E/A) and the mutant lacking the cytoplasmic domain (ΔCyto). Tfap2α.L-S1 and Tfap2α.L-S3 were cloned from xenopus embryos using Takara infusion cloning. The Arid3a clones was described elsewhere ^6^. The luciferase promoters were all cloned into pGl3 (Adgene). The genomic sequence for *Xenopus laevis* promoters were cloned based on their position in the genome. The Tfap2α−S1 promoter correspond to 2368 bp from the first ATG, the Tfap2α−S3 promoter correspond to 3577 bp from the third ATG and does not overlap with the S1 promoter. The *Calpain-8.L* promoter correspond to 2483 bp upstream of the first ATG. Human Arid3a shRNA was bought from Sigma-aldrich (TRCN0000013791, target sequence CCCTAAGATCAAGAAAGAGGA).

### Injections and microdissections and mRNA synthesis

SP6 polymerase was used for capped mRNAs synthesis after digesting using Not1 ^13^. Injectors were calibrated using a 1 µL capillary needle (Microcaps, Drumond, PA, United States). The injection pressure was set at 15 psi and the injection time set between 50 and 200 ms to obtain a 5 nL delivery. Embryos were injected at 1-cell, 2-cell, 8-cell and 16-cells as described previously ^53^. Embryos were raised at 14°C–15°C until scoring for neural tube closure or CNC cell migration. For each injection, the percentage of fluorescent CNC present in the branchial pathway was normalized to control embryos and was set to 100%.

### Generation of Knockout Adam13 Line

Targets in Exon 1 and Exon 5 of *adam13.L* were identified using CRISPRScan (https://www.crisprscan.org/) ^68^ and the second nucleotide of each Single guide RNA was modified to a G at the 5’end, for increased mutagenic activity ^69^, T1: GGGCACATGGCTGGGACTCG and T2: GGAGGTGGTGAGGGCGACAG. Single guide RNAs were synthesized by in vitro transcription of the sgRNA PCR template using the T7 MEGAscript kit (Ambion, Cat. No. AM1334). To generate eggs, inbred *Xla.J-Strain^NXR^ (RRID:* NXR_0024) Xenopus laevis females were first given 35 U Pregnant Mare Serum Gonadotropin (PMSG) (Bio Vender, Cat. No. RP17827210000) followed by 350 U of Human Chorionic Gonadotropin (hCG) (Bio Vender, Cat. No. RP17825010) to induce ovulation. Eggs were then fertilized using methods previously outlined in Shaidani et al. ^70^. Founders were generated by coinjecting 250pg of sgRNA 1 and 250pg of sgRNA 2 with 500pg of Cas9 protein per embryo at the one cell stage. Embryos were collected and genotyped using Qiagen DNeasy Blood and Tissue (Cat. No / ID: 69506). Genomic DNA was amplified using the following PCR primers: Exon 1 forward primer 5’-GTGTTGGGTGAGTTTATTGGGC-3’ and reverse primer 5’-TGATAGAGCCAGGCTATCAGGG-3’; Exon 5 forward primer 5’-CTTCGGACATTGTGATCCTTGC-3’ and reverse primer 5’-TTGATGAGATATAGTCGCCCGG-3’. Embryos were obtained from seven F0 *adam13* knockout males by outcrossing to J-strain females. F1 embryos were then genotyped to determine if germline mutations were present. Of the seven males, only three had germline mutations in exon 1 (Male 2: -13/+, Male 6: -7/+, Male 7: -13/+). No mutations were found in exon 5. Offspring from Male 2 were then raised through metamorphosis and intercrossed to generate -13/-13 *adam13* Knockouts. This mutation results in a premature stop following amino acid 22. The -13/-13 (*Xla.adam13^emNXR^*, RRID: NXR_2117) *adam13* mutants are available from the NXR.

### Cell culture and transfection

HEK293T (ATCC, CRL-3216 RCB2202) cells were cultured in RPMI media supplemented with 10 U/mL Pen/Strep, 2 mM L-glutamine, 0.11 mg/mL sodium pyruvate, and 10%FBS (Hyclone, South Logan, UT). Transfections were performed using Polyethylenimine (PEI, Polysciences Inc.). For each transfection, 1 µg of DNA was mixed with 10 µg of PEI in 200 µL of RPMI media (Hyclone) at room temperature for 15 min prior to adding to the cells dropwise. The media was changed 24 h post transfection.

### Immunoprecipitation and western blots

HEK293T, embryos and CNC explants were extracted in 1XMBS-1% Triton-X100, Protease phosphatase inhibitor cocktail 1X (Thermoscientific) and 5 mM EDTA. Immunoprecipitations were performed using 1-5 ug of antibody bound to protein A/G magnetic beads (Thermofisher), incubated overnight at 4°C. The beads were washed 3 times for 5 min with extraction buffer at room temperature then eluted in 2X reducing laemmli sample buffer. All proteins were separated in 5%–22% gradient SDS-PAGE gels and transferred to polyvinylidene fluoride membranes (PVDF, Millipore, Billerica, MA) using a semi-dry transfer apparatus (Hoeffer).

### Immunofluorescence

CNC cells were dissected at stage 17 and placed on fibronectin coated glass bottom plates (20ug/mL) for 3 h at 18°C ^71,72^. The explants were fixed using MEMFA (0.1 M MOPS pH 7.4, 2 mM EGTA pH 8, 1 mM MgSO4 and 4% paraformaldehyde) for 1 h, permeabilized using 0.5% TX100 in 1XMBS with 100 mM glycine for 1 h. The explants were then blocked using PBS containing 10% heat inactivated goat serum, 1% BSA, 0.1% Tween for 1 h prior to incubation in the same buffer with the primary antibodies overnight at 4°C. CNC explants were washed in PTw (PBS-tween 0.1%) 3 times 15 min, blocked again for 1 h at room temperature using blocking solution prior to incubation with the secondary antibody and Hoest33342 for 1 h at room temperature. The explants were washed 3 times in PTw and imaged using Nikon confocal microscope (A1RHD25).

### Immunofluorescence analysis

The images were taken at the same gain and power for all explants within the same experiment. The Immunofluorescence images were analyzed using GA3 Nikon. The intensity in the nucleus was measured by making a mask using the marker of CNC Sox9 or Snai2, then the intensity of the target was normalized with the intensity of Hoest33342. T-test was used to determine significance in at least 3 independent experiment.

### Mass spectrometry

Protein extracted and immunoprecipitated with the target-specific antibody from embryos or HEK293T cells were resuspended in 8M urea and processed with Trypsin/Lysine-C according to manufacturer instruction (Promega). All samples were cleaned using Zip-Tip (Fisher Scientific). For BioID experiments HEK293T cells were transfected with Adam13-BioID tagged constructs, and grown in media depleted for Biotin (Streptavidin depletion). Biotin was added to the media at various time after transfection. Transfected cells were either directly extracted as described above or cytoplasmic and nuclear extraction was performed (Pierce NE-PER kit). The Biotinylated proteins were then purified using streptavidin magnetic beads (Pierce) before denaturation in 8M urea and processing with Trypsin/Lysine-C according to manufacturer instruction (Promega). Tandem mass spectrometry was performed using a Thermo orbitrap Fusion (Mass spectral data were obtained at the University of Massachusetts Mass Spectrometry Core Facility, RRID:SCR_019063). All MS/MS samples were analyzed using Sequest (Thermo Fisher Scientific, San Jose, CA, United States; version IseNode in Proteome Discoverer 2.4.1.15). Sequest was set up to search a Xenopus protein database produced from the Xenopus laevis genome (Xenbase 72,266 entries) or the human protein database uniprot-human containing the sequences for Xenopus Adam13, Arid3a and Tfapα (126,358 entries) assuming the digestion enzyme trypsin. Sequest was searched with a fragment ion mass tolerance of 0.60 Da and a parent ion tolerance of 10.0 PPM. Carbamidomethyl of cysteine was specified in Sequest as a fixed modification. Oxidation of methionine, acetyl of the n-terminus and phospho of serine threonine and tyrosine were specified in Sequest as variable modifications.

Criteria for protein identification: Scaffold (version Scaffold_5.0.1, Proteome Software Inc., Portland, OR) was used to validate MS/MS based peptide and protein identifications. Peptide identifications were accepted if they could be established at greater than 95.0% probability by the Peptide Prophet algorithm ^73^ with Scaffold delta-mass correction. Protein identifications were accepted if they could be established at greater than 99.0% probability and contained at least 2 unique identified peptides. Protein probabilities were assigned by the Protein Prophet algorithm^74^. Proteins that contained similar peptides and could not be differentiated based on MS/MS analysis alone were grouped to satisfy the principles of parsimony. Proteins sharing significant peptide evidence were grouped into clusters.

### RNA-sequencing and analysis

RNA was prepared using Roche RNA extraction kit (11828665001) from dissected CNC. Isolated RNA was sent for sequencing to Azenta life sciences to perform paired end long-read sequencing (2X150,∼60M reads). Sequenced RNA was trimmed for adapters and low-quality reads using trimmomatic ^75^. The resulting reads were mapped to Xenopus laevis genome v10.1, using Spliced Transcripts Alignment to a Reference (STAR) ^76^. The mapped reads were counted for genes as well exon specific counts using feature counts in the Subread package ^77^. Differences in fold changes between the conditions was analyzed using DESeq2 ^78^. For pathway analysis and heatmap preparation, the gene names were converted into Human orthologs. The pathway analysis was performed using GSEApy ^79^ and GSEA ^44^ with clusterprofiler ^80^. Sashimi plots were made using Integrative Genomics Viewer (IGV) ^81,82^. Custom heatmaps were made using geneset from GSEA.

### Statistical analysis

If not specified in the figure legends, for two samples two-tail unpaired Student’s t-test was performed. For more than 2 samples one-way ANOVA was performed. The analysis was performed in the software Graphpad-Prism.

## Figure Legends

**SUPPLEMENTARY FIGURE S1.**
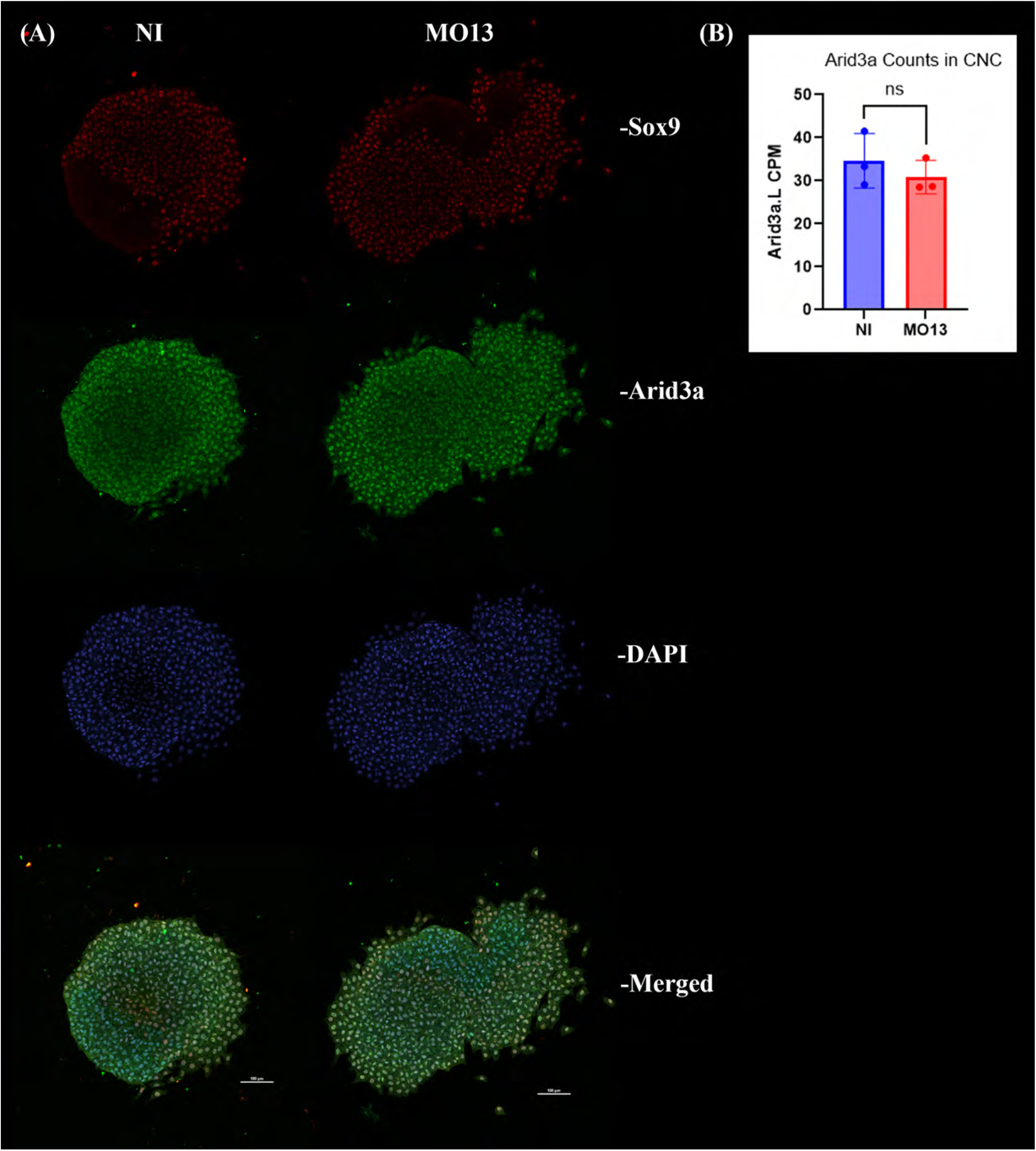
**(A) Immunofluorescence images of CNC** explants dissected at stage 17 and grown on fibronectin substrate for 3hrs. CNC explants were fixed and subsequently immuno-stained for Sox9 (red), Arid3a (green) and DAPI (blue) and imaged using confocal microscopy; all images are Max-IP. (B) Histogram depicting CPM (counts per million) of Arid3a.L RNA present in CNC RNA-sequencing. Student t-test was performed for statistical analysis, ns represents p-value>0.05.

**SUPPLEMENTARY FIGURE S2.**
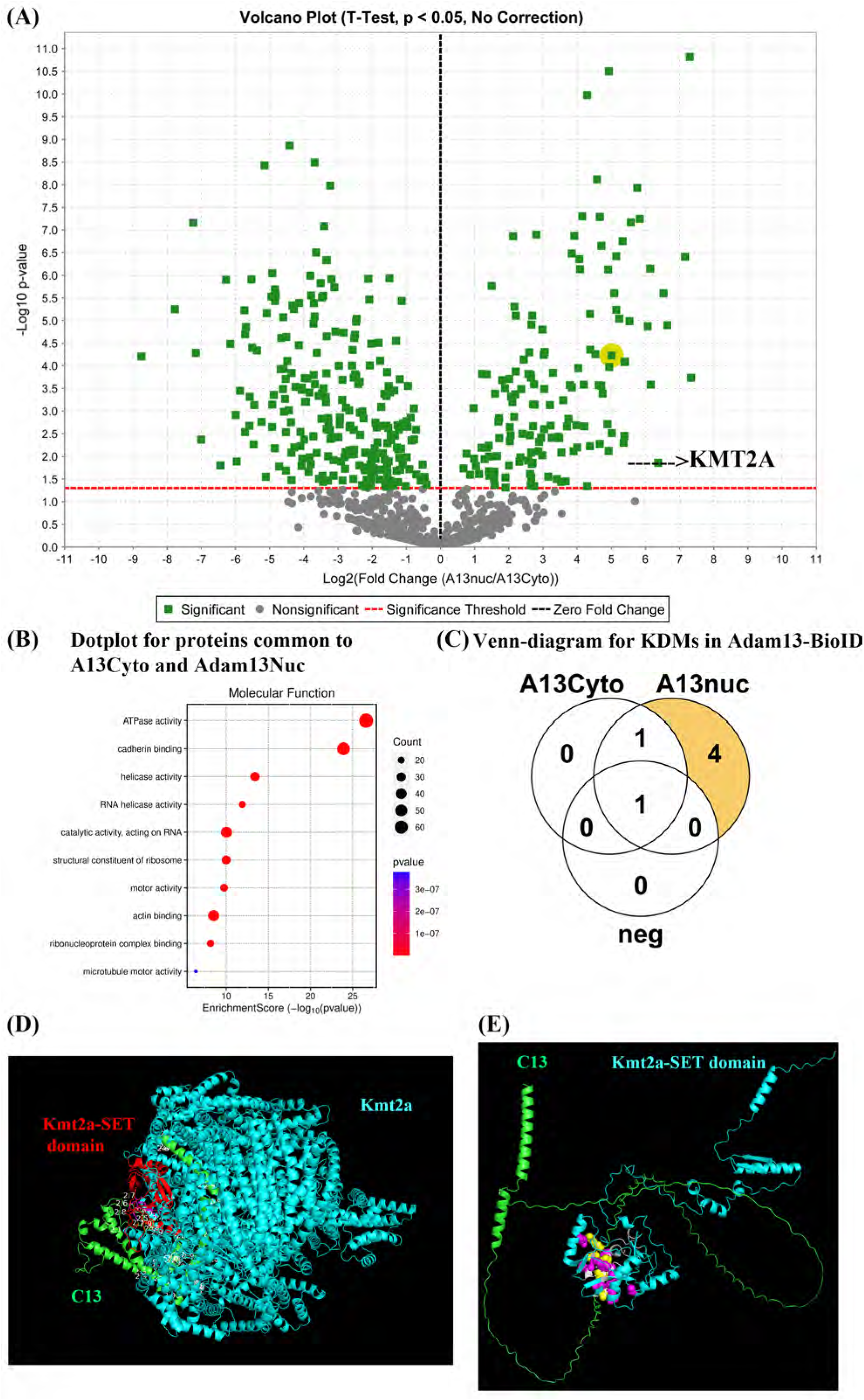
(A) Volcano plot representing proteins detected in the mass spectrometry of proteins labelled by Adam13-BioID. Squares in green represent proteins found statistically significant between the Adam13 cytoplasmic fraction (A13Cyto, 6 samples) and Adam13 nuclear fraction (A13nuc 6 samples). Proteins identified from cells that were transfected with constructs lacking the Bio-ID tagged (6 samples) were removed prior to the analysis. The red line corresponds to the minimum significant value. The right upper quadrant represents proteins significantly enriched in the nuclear fraction, the left quadrant represents those enriched in the cytoplasmic fraction. The green square highlighted in yellow is KMT2A enriched in the nuclear fraction. (B) Dot plot representing the gene set enrichment pathway analysis for molecular function of proteins that were found biotinylated in both cytoplasmic and nuclear fraction. The red color represents higher significance (lower p-value), and the size of the dot represents the number of genes in the set. (C) Venn diagram representing total KDM proteins detected at 2 unique-peptide minimum for Control (neg), Adam13 cytoplasmic fraction (A13Cyto), and Adam13 nuclear fraction (A13nuc). (D) Alphafold3 model of the interaction between the cytoplasmic domain of Adam13 (green) and Kmt2a (Cyan). The Set domain of Kmt2a is represented in red. (E) Aplhafold3 model of the interaction between the cytoplasmic domain of Adam13 (Green) and the Set domain of Kmt2a (Cyan). Residues predicted to interact (3A) in both chains are highlighted (Purple for Set and Yellow for Adam13).

**SUPPLEMENTARY FIGURE S3.**
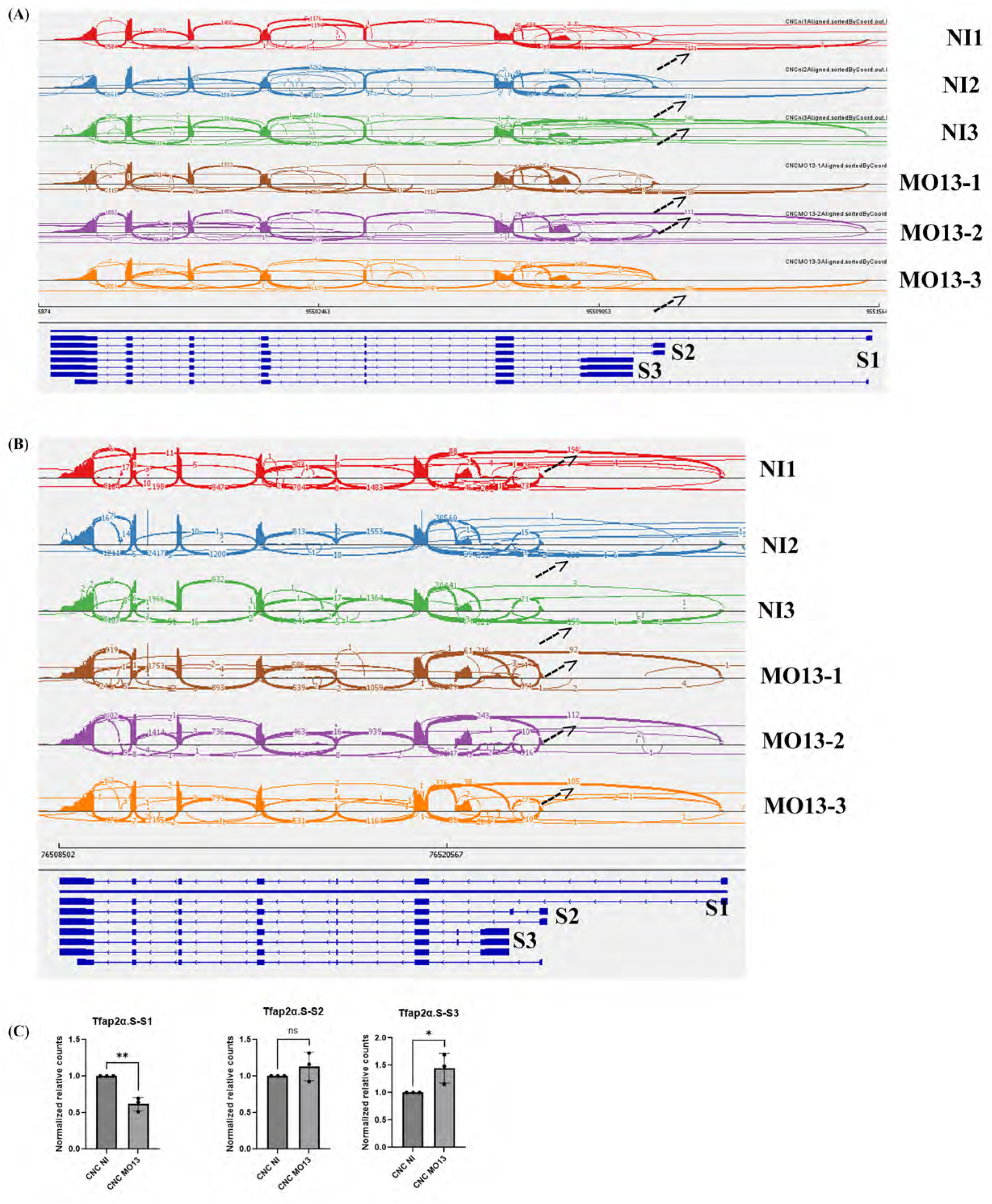
(A) Sashimi plot analysis of transcript junctions detected for gene Tfap2α.L in RNA-sequencing data between Control (NI1, NI2, NI3) and Adam13KD (MO13-1, MO13-2, MO13-3), numbers on the line are the major transcript junction between corresponding exons, arrow points to the number of transcript junction detected between Start1 and Exon2. (B) Sashimi plot analysis of transcript junctions detected for gene Tfap2α.S in RNA-sequencing data between Control (NI1, NI2, NI3) and Adam13KD (MO13-1, MO13-2, MO13-3), numbers on the line are the major transcript junction between corresponding exons, arrow points to the number of transcript junction detected between Start1 and Exon2. (C) Quantification of exon specific counts for Tfap2α.S-start1, start2 and start3 normalized to total transcript counts for Tfap2α.S gene. Student t-test was performed for statistical analysis, * represents p-value<0.05, ** represents p-value<0.01, ns represents p-value>0.05.

**SUPPLEMENTARY FIGURE S4.**
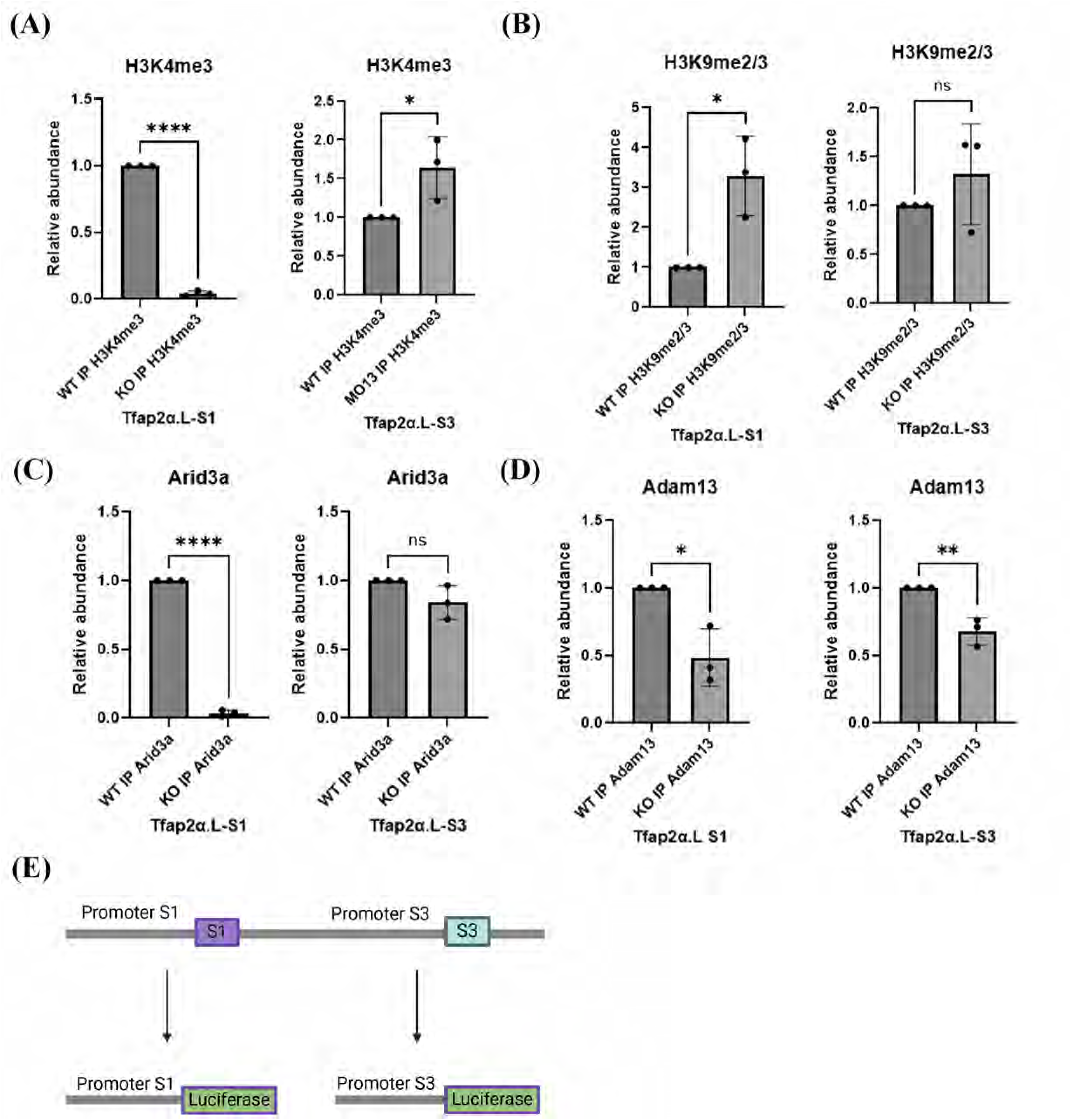
Chromatin-Immunoprecipitation-qPCR analysis of DNA associated with the chromatin binding proteins. (A, B, C, D) Histogram depicting the relative abundance of DNA bound to H3K4me3 (A), H3K9me2/3 (B), Arid3a (C), Adam13 (D) in WT and KO embryos at the promoter region of Tfap2α-S1 and Tfap2α-S3 based on ChIP-qPCR analysis. ChIP was performed at stage 20 (Independent biological replicate from figure 4). Student’s t-test was performed for statistical analysis, * represents p-value<0.05, ** represents p-value<0.01, *** represents p-value<0.001, **** represents p-value<0.0001, ns represents p-value>0.05. (E) Schematic representation of the Luciferase reporters for the S1 and S3 transcription starts.

**SUPPLEMENTARY FIGURE S5.**
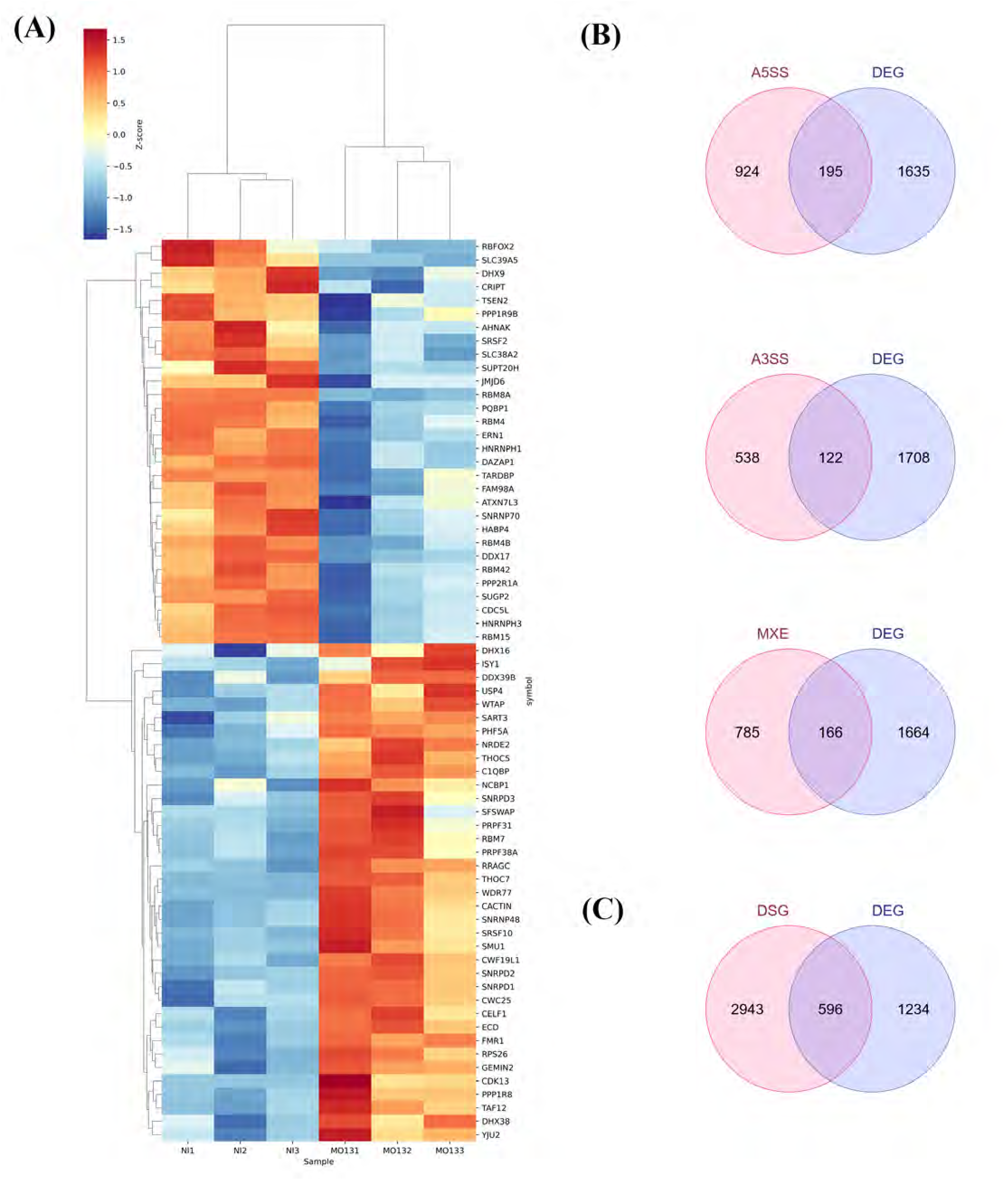
(A) Custom heatmap visualization of genes derived from comparing Human Gene Set: REACTOME_MRNA_SPLICING with differentially expressed genes in CNC RNA-sequencing data between Control (NI1, NI2, NI3) and Adam13KD (MO131, MO132, MO133), higher Z-score in red representing higher expression and blue representing lower expression. (B) Venn diagram depicting the genes found differentially expressed (p-value<0.05) versus genes having Alternate 5’ splice site (A5SS), Alternate 3’ splice site (A3SS), and mutually exclusive exons (MXE). (C) Venn diagram representing, differentially spliced (DSG) to differentially expressed (DEG) genes.

**SUPPLEMENTARY TABLE S1.** Proteins labelled by Adam13-BioID in the cytoplasm and the nuclear fractions (6 biological repeats each, minimum of 2 exclusive unique peptides). Proteins that were found in sample lacking BioID have been removed. (A13Cyto) represent proteins significantly more associated with Adam13 in the cytoplasm (p<0.05). (A13Nuc) represent proteins significantly more associated with Adam13 in the nucleus (p<0.05). (Common) represent proteins that were found in both samples.

**SUPPLEMENTARY TABLE S2.** List of genes significantly affected by the loss of Adam13 in the CNC. Log fold change, pvalue and adjusted pvalues are provided for each gene.

**SUPPLEMENTARY TABLE S3.** List of genes significantly affected by the loss of Adam13 in the CNC. Upregulated and Down regulated genes are indicated in two lists (Up yellow, down blue). The description of the Human orthologue function is presented.

**SUPPLEMENTARY TABLE S4.** List of proteins associated with the different Tfap2α variants (minimum of 2 exclusive unique peptides). Any proteins found in the Flag IP from the negative control (RFP-Flag) was eliminated. Proteins that were significantly associated with either variant are indicated in separate lists (TFAP2A-S1 or S3). The proteins that were found associated with both variants are in the COMMON list.

**SUPPLEMENTARY TABLE S5.** List of genes differentially expressed and differentially spliced in CNC lacking Adam13. Differentially expressed (DEG), alternate 5’ splice site (A5SS), alternate 3’ splice site (A3SS), and mutually exclusive exons (MXE).

**SUPPLEMENTARY TABLE S6.** List of proteins identified with 2 minimum exclusive unique peptides in human ADAM9, Xenopus Adam13 Immunoprecipitation and Adam13-BioID experiments. Proteins identified in any of the negative control were eliminated.

**SUPPLEMENTARY TABLE S7.** List of proteins found in wild type Adam13 and Adam13-833D but not Adam13-833A or Adam13ΔCyto immunoprecipitation are provided. The String representation of interaction for these proteins is also provided. Proteins involved in mRNA transport (Red) and mRNA splicing via the spliceosome (Blue) are also indicated.

